# EEG connectome-based predictive modeling of nonverbal intelligence level in healthy subjects

**DOI:** 10.1101/2024.11.04.621793

**Authors:** Anton Pashkov, Ivan Dakhtin, Inna Feklicheva, Julia Shmotina, Mahmoud Hassan

**Affiliations:** FSBI “Federal Center of Neurosurgery“, Novosibirsk, Russia; Department of neurosurgery, Novosibirsk State Medical University, Novosibirsk, Russia; Department of Data Collection and Processing Systems, Novosibirsk State Technical University, Novosibirsk, Russia; School of Medical Biology, South Ural State University, Chelyabinsk, Russia; Department of Fundamental Medicine, Chelyabinsk State University, Chelyabinsk, Russia; MINDIG, Rennes, France; School of Science and Engineering, Reykjavik University, Reykjavik, Iceland

**Keywords:** EEG, intelligence, resting state, connectome, machine learning

## Abstract

Intelligence is increasingly recognized as a critical factor in successful behavioral and emotional regulation. Neuroimaging techniques coupled with machine learning algorithms have proven to be valuable tools for uncovering the neural foundations of individual cognitive abilities. Nevertheless, current electroencephalograph (EEG) studies primarily focus on classification tasks to predict the intelligence category of subjects (e.g., high, medium, or low intelligence), rather than providing quantitative intelligence level forecasts. Furthermore, the outcomes obtained are significantly impacted by the specific data processing pipeline chosen, which could potentially compromise result generalizability. In this study, we implemented a connectome-based predictive modeling approach on high-density resting state EEG data from healthy participants to predict their nonverbal intelligence level. This method was applied to three independently collected datasets (N = 255) with different functional connectivity methods, parcellation atlases, threshold p-values and curve fitting orders used to ensure the reliability of the findings. We found that the prediction accuracy expressed in terms of R² varied significantly depending on the processing pipeline configuration, ranging from negative R^2^ values up to 0.27. The most consistent results across datasets were found in the alpha frequency band. Furthermore, we employed a computational lesioning approach to identify the valuable edges that made the most significant contribution to predicting intelligence. This analysis highlighted the crucial role of frontal and parietal regions in complex cognitive computations. Overall, these findings support and expand upon previous research, underscoring the close relationship between alpha rhythm characteristics and cognitive functions and emphasizing the critical consideration of method selection in result evaluation.

## Introduction

Intelligence stands as a fundamental and extensively studied concept within contemporary psychology, intricately connected to a wide array of cognitive abilities exhibited by humans. Its influence extends to various aspects of individuals’ lives, including educational attainment (Rask-Andersen et al., 2021), occupational achievement (Plomin & Deary, 2015), overall quality of life (Geary, 2019), and psychosocial adaptation (Huepe et al., 2011). Moreover, individuals with higher intelligence possess a more adaptable and diverse repertoire of cognitive resources known as cognitive reserve, which enables them to effectively adjust problem-solving strategies, optimizing approaches without compromising solution efficiency (Whalley et al., 2016). In the structure of general intelligence, nonverbal abilities refer to “the ability to represent, transform, generate, and recall symbolic, non-linguistic information”, elevated proficiency in which also dictates accomplishments in disciplines like science, technology, engineering, and mathematics (Linn & Petersen, 1985), (Shakeshaft et al., 2016; Uttal et al., 2013).

As is the case with nearly all complex phenomena under investigation, research on intelligence, including nonverbal abilities, led to a great variety of ways in which one can define, classify and gauge it. Among the most widely used tools for intelligence measurement are Wechsler Adult Intelligence Scale (WAIS), Raven progressive matrices (RPM), Penn Matrix Reasoning Test (PMAT) and, created at the dawn of intelligence measurement era, Stanford-Binet Intelligence Scale, all showing good to excellent psychometric characteristics. Another available option is to derive a general factor of intelligence (g) based on scores taken from a set of cognitive tasks (Gläscher et al., 2010).

However, the evaluation of intelligence through conventional paper-and-pencil tests or their computerized counterparts presents several drawbacks. The availability of most intelligence tests impacts the reliability of cognitive function assessment. Moreover, intelligence measurement via tests heavily relies on the test content, which may become outdated over time, and the creation of new tests necessitates substantial time investment. Currently, efforts are underway to develop methodologies for intelligence assessment that employ machine learning techniques, leveraging task-based or resting-state neuroimaging data. These approaches aim to account for the complex interplay between the level of cognitive ability development and the neurobiological mechanisms that underlie individual variations in intelligence (Choi et al., 2008; Hilger et al., 2017b; Pietschnig et al., 2015).

For an extended period, magnetic resonance imaging (MRI) and functional MRI have been the dominant noninvasive neuroimaging techniques in this research field, offering scientists valuable insights into the complex relationship between human mental abilities and patterns of neural activity. Notably, fMRI studies have allowed researchers to shift their focus from examining static aspects of brain organization to dynamic ones. The task-based paradigm in neuroimaging has been successful in identifying key brain regions that potentially form the neural foundation for cognitive computations. Specifically, the frontal and parietal cortices have emerged as pivotal in this regard (Cole et al., 2012; Jung & Haier, 2007).

Subsequent investigations that adopted a resting state perspective revealed that these approaches were equally informative and productive in uncovering new insights into the neural underpinnings of intelligence compared to task-based studies (Li et al., 2018). A multitude of studies utilizing fMRI to examine brain network dynamics have demonstrated that intelligence level is tightly linked to both global and local properties of brain functional connectivity (Hilger et al., 2017a; Kruschwitz et al., 2018).

Despite its overall success, the utilization of MRI methods encounters several obstacles. MRI equipment is costly, cumbersome, and necessitates a specially shielded room for data acquisition, thereby impeding its widespread and convenient usage. Furthermore, a notable limitation of fMRI studies is the relatively poor temporal resolution (2-3 seconds), which is constrained by the pace of hemodynamic response, while cognitive processes occur at a faster rate (Amaro & Barker, 2006). In stark contrast, electroencephalography (EEG), another noninvasive neuroimaging technique, circumvents the aforementioned drawbacks inherent to MRI. Presently, EEG is regarded as one of the most dependable and informative diagnostic tools in the field of neuroscience. The scientific and technological advancements in recent decades have greatly enhanced the recording technology and methods for EEG analysis (Michel & Murray, 2012). Notably, the utilization of high-density EEG, which permits the recording of neuronal activity from 64, 128, and 256 channels, is gaining popularity. A significant milestone has been achieved in addressing the inverse EEG problem - the localization methods for source estimation implemented in specialized software have been successfully validated through joint EEG-fMRI registration and invasive electrocorticography in patients undergoing neurosurgery (Cohen, 2017).

One of the advantages of employing EEG is its ability to not only identify the brain regions involved in cognitive processes but also discern the frequency characteristics of their interaction. Notably, EEG can also reconstruct macroscale networks, similar to those derived from fMRI, thereby enabling scientists to test hypotheses generated from fMRI findings using EEG (Wirsich et al., 2021; Yuan et al., 2016).

The number of studies conducted utilizing EEG is considerably lower compared to those employing MRI. However, despite this disparity, the findings from EEG studies, which partially replicate and partially complement the results obtained from MRI, merit significant attention and consideration (Nentwich et al., 2020; Wirsich et al., 2021).

Various EEG characteristics have been employed to investigate the relationship between intelligence and patterns of neural activity. Langer and co-authors discovered that nonverbal intelligence exhibited associations with functional connectivity values derived from EEG data recorded during a resting state. Specifically, significant correlations were observed between Raven Progressive Matrix scores and graph metrics of alpha band brain networks: positive correlations were found for the Small World Index and clustering coefficient, while a negative correlation was observed for the characteristic path length (Langer et al., 2012). In another recent study, Zakharov and colleagues found that the characteristics of path length within the alpha band range in brain networks were significantly correlated with non-verbal intelligence in the sensor space, but not in the source space (Zakharov et al., 2020).

The integration of neuroimaging techniques and machine/deep learning methods is rapidly gaining traction in the field of neuroscience. This combination is being utilized to address a diverse array of tasks when working with large-scale datasets. These tasks range from predicting biological age, cognitive functioning, and emotional responses to identifying specific patient groups affected by particular diseases (Nielsen et al., 2020; Niu et al., 2020; Yang et al., 2023). Notably, the forefront of this trend lies in the prediction of interindividual variations in IQ using machine/deep learning methods applied to neuroimaging data.

In this paper, our focus was specifically on one machine learning method that has demonstrated its effectiveness in addressing such challenges within the past five years - the connectome-based predictive modeling framework (CPM). CPM is a data-driven approach that utilizes multivariate regression to forecast individual differences in cognitive abilities and behavioral traits based on connectome data (Shen et al., 2017). CPM has been successfully applied in various domains, including psychiatry, neurology, and psychology, it has predominantly been employed with functional magnetic resonance imaging (fMRI) data, with only a few exceptions (Boyle et al., 2023; Kabbara et al., 2022; Wang et al., 2021; Wu et al., 2023; Yoo et al., 2018).

The initial application of the predictive framework was conducted in a study by Finn et al., wherein they discovered that fluid intelligence, as estimated from Progressive Matrices, could be forecasted using functional connectivity matrices within a relatively small sample size (n = 118) from the Human Connectome Project (HCP) dataset. The correlation between observed and predicted scores was reported as r = 0.5. Subsequently, the authors revised the effect size to r = 0.2 based on a larger dataset (n = 606) (Finn et al., 2015; Noble et al., 2017).

Furthermore, in a separate study, Dubois et al. employed the predictive framework on a sample of 884 individuals from the HCP database. They successfully predicted approximately 20% of the variance in general intelligence using participants’ resting-state connectivity matrices (though the effect size was later revised to approximately r=0.09). This prediction was not dependent on any specific anatomical structure or network, but rather on redundant information dispersed throughout the brain (Dubois et al., 2018).

The vast majority of studies exploring the relationship between brain and intelligence have primarily utilized fMRI techniques. However, it is worth noting a few noteworthy studies that have employed EEG as well. For instance, Nentwitch et al. utilized multivariate distance matrix regression to establish a connection between various behavioral phenotypes and connectivity patterns. They found a significant association between IQ and EEG activity specifically within the gamma and beta frequency ranges (Nentwich et al., 2020). Another notable study by Friedman et al. involved participants solving Raven’s Matrices at progressively increasing levels of complexity during an EEG recording session. They defined cognitive load based on problem difficulty and successfully demonstrated that it could be predicted from the EEG readings of the participants (Friedman et al., 2019). However, it is important to note that, to the best of our knowledge, no study has been conducted thus far that enables a quantitative prediction of intelligence levels in healthy individuals using resting-state EEG data.

In this article, we utilize connectome-based predictive modeling applied to high-density resting-state EEG data to predict levels of nonverbal intelligence in a cohort of healthy subjects. Our study is distinct as it is the first one to quantitatively predict nonverbal intelligence levels using EEG connectome data. Importantly, the subjects were drawn from three independent samples collected by different research teams. Furthermore, to ensure the reliability and robustness of our findings, we employed different data processing pipelines, systematically varying factors such as brain parcellation schemes, functional connectivity methods, and p-value thresholds. By doing so, we aimed to assess the impact of these confounding factors on the predictive performance of the CPM approach. Additionally, we explored the implementation of non-linear variants of CPM to examine whether such modifications could enhance the predictive ability of the model. These rigorous analyses aimed to strengthen the validity and generalizability of our results.

## Materials and methods

### Experimental design

The study utilized three resting-state EEG datasets, which are described in detail below. These datasets underwent identical processing steps outlined in Figure 1. Initially, the data were preprocessed to remove artifacts before source reconstruction. Subsequently, the source time series were analyzed to evaluate their pairwise interactions and generate functional connectivity matrices. These matrices were then fed into the CPM algorithm to predict intelligence scores. CPM was executed for each participant using an internal leave-one-out cross-validation approach (LOOCV). Lastly, external cross-sample validation was applied to validate the results.

**Figure 1.**
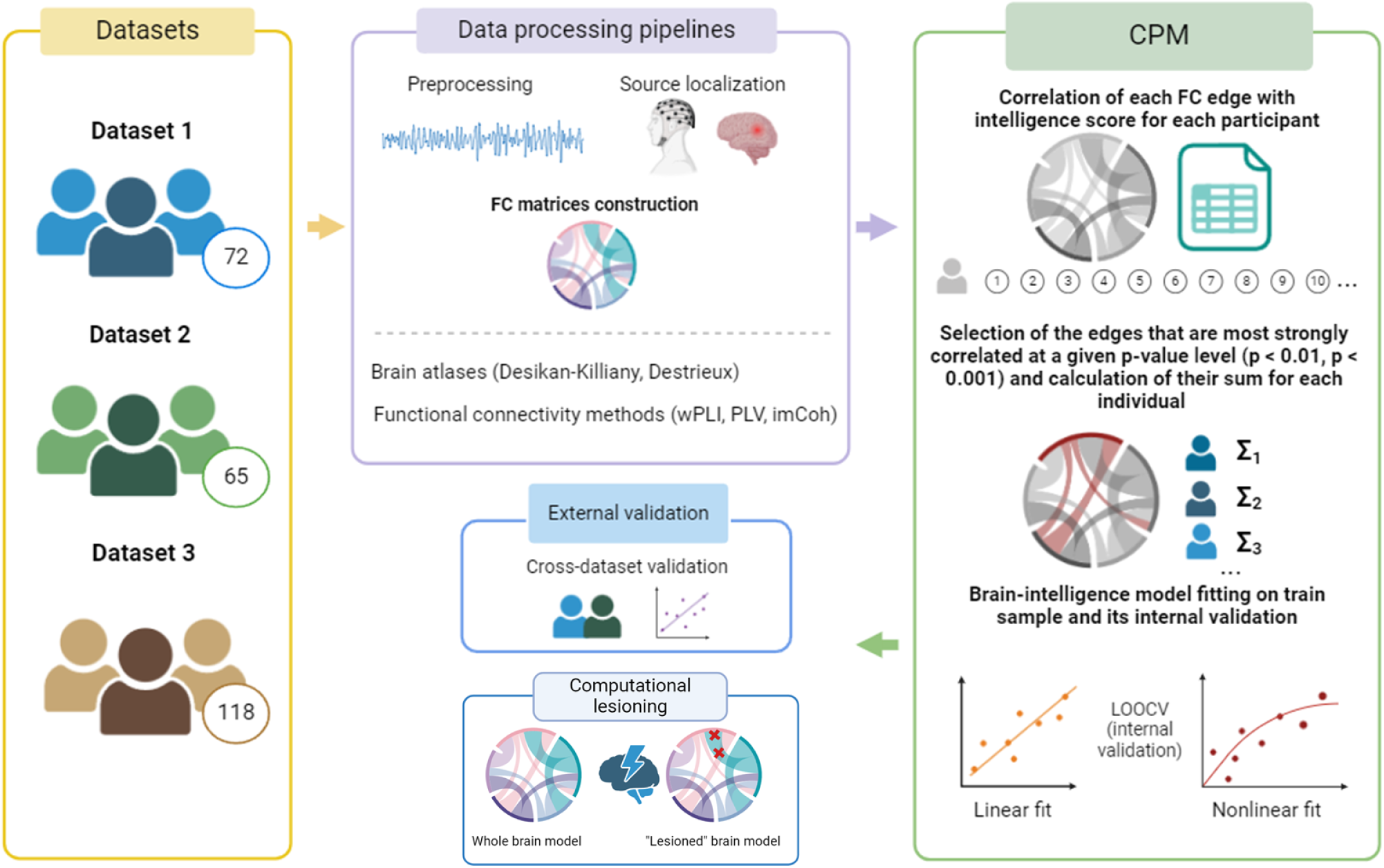
General design of the study.

**Figure 2.**
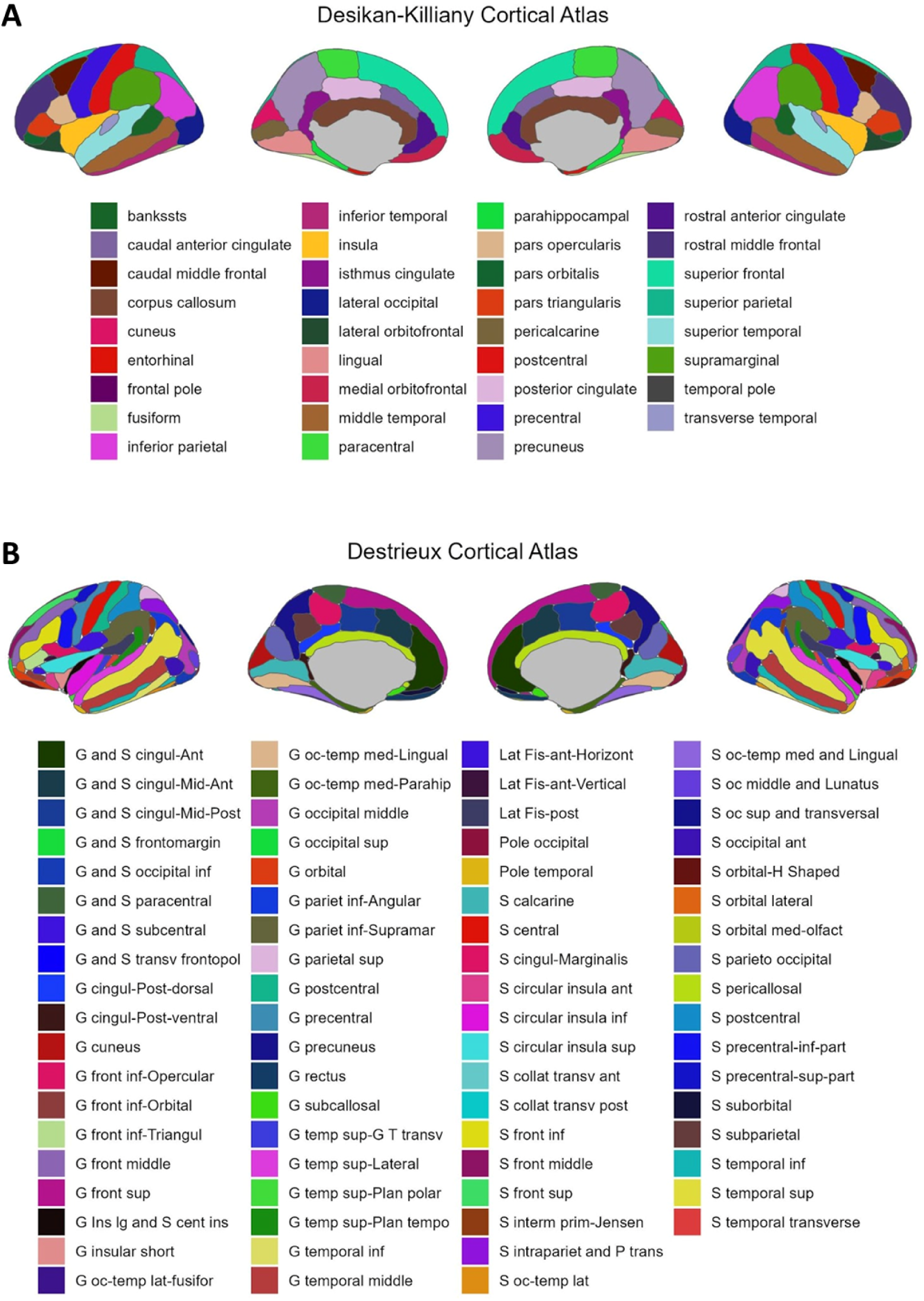
Two brain parcellation atlases used in the study. Panel A: Desikan-Killiany cortical atlas. Panel B: Destrieux cortical atlas.

In this study, we focused on a specific set of parameters as the baseline configuration. We chose the Destrieux atlas over the DK atlas for its finer granularity. Setting the p-value threshold to 0.001 allowed us to reduce false positives more effectively. Additionally, we utilized partial correlations within the CPM procedure to mitigate the impact of age on intelligence. To evaluate how these parameters could affect the predictive accuracy, we explored alternative configurations by varying one parameter at a time. These variations included adjusting the p-value threshold to 0.01, changing the atlas to Desikan-Killiany, and modifying the orders of CPM to 1, 2, or 3.

### Datasets

We utilized three datasets, each consisting of 62 or 64-channel EEG (depending on the particular dataset) recordings and intelligence measures, with a total of 255 participants included.

#### Dataset # 1

In the first sample, data of 72 participants (41 females) aged 18 to 25 years old (median age: 20 years old) were collected in the Molecular and Genetic Research Lab at South Ural State University, Chelyabinsk. The study was approved by the ethical committee of South Ural State University and conducted in accordance with relevant ethical guidelines (Declaration of Helsinki). Prior to data acquisition, all participants provided written informed consent. Participants were recruited from introductory psychology classes and received course credits as compensation for their participation. Recruitment criteria included having no history of head trauma or seizures, abstaining from psychoactive substances including alcohol and caffeine for at least one day before EEG recording session, having no psychiatric or neurological disorders, and having slept for a minimum duration of 6 hours on the previous night. During the EEG resting state session, participants were instructed to relax for ten minutes while alternating between open eyes and closed eyes states every two minutes. EEG data was recorded using a Brain Products ActiChamp amplifier with 64 electrodes placed according to the international 10-10 system. The experiments took place in a soundproof electrically shielded room with dim lighting. Impedance levels were maintained below 25 kOhm using highly conductive chloride gel. Continuous EEG recordings without any filtering or downsampling were performed using Brain Products PyCorder acquisition system at a sampling rate of 500 Hz; the referencing electrode was located at point Cz (re-referenced to average). The recorded data was downsampled to 200 Hz and bandpass filtered from 0.1 Hz to 45 Hz range. Markers indicating closed and open eyes conditions were added to separate continuous EEG fragments recorded during respective conditions. To estimate nonverbal cognitive abilities of participants, Raven’s Standard Progressive Matrices (RSPM) test consisting of series-based items (A → B → C → D → E) was administered, with increasing complexity from series to series. Participants were given a maximum of 20 minutes to complete the test, and the total number of correct answers served as a general indicator of nonverbal abilities.

#### Dataset # 2

The second sample collected during the execution of Cuban Human Brain Mapping Project (Valdes-Sosa et al., 2021). This study followed the Declaration of Helsinki and was approved by the Ethical Review Committee of the Cuban Neuroscience Center. Exclusion criteria were the presence of any somatic disease or brain dysfunctions. Resting-state EEG data were recorded using MEDICID 5 digital EEG system with either 64 or 128 electrodes. For the analysis, we included data from 65 participants (54 males, 11 females) recorded using a 64-channel device, with participants’ ages restricted to between 18 and 35 years old (median age of 26). The EEG recordings lasted at least 30 minutes and encompassed conditions such as eyes closed, eyes open, hyperventilation, and subsequent recovery. Electrodes were placed according to the international standard “10-10” system using a customized electrode cap; linked earlobes served as reference for EEG signals. Electrode impedances below 5 KΩ were considered acceptable for recording quality assurance purposes. The bandpass filter parameters were set between frequencies of 0.5 Hz to 50 Hz with an additional notch filter at frequency of 60 Hz. All signals were sampled at a rate of 200 Hz. The EEG recordings took place in a controlled environment that maintained stable temperature levels while participants sat on reclined chairs instructed to relax and remain still throughout testing in order to minimize movement-related artifacts and excessive blinking. In addition to EEG measurements, a battery of behavioral measures included MMSE scale, WAIS-III, and Go No-go test. Intelligence levels were assessed using the fully validated and translated Spanish version of the David Wechsler Adult Intelligence Scale (WAIS-III). The scale provided scores for Full-Scale IQ (FSIQ), Verbal IQ (VIQ), Performance IQ (PIQ). The raw intelligence measures were scored according to official normative data from the printed WAIS-III version. However, to mitigate cultural bias, the scores were subsequently standardized based on information derived from the Cuban sample to generate age-adjusted performance scores specific to the population.

#### Dataset # 3

The third sample was obtained from MPI (Max Planck Institut) Leipzig Mind-Brain-Body Dataset (Babayan et al., 2019; Mendes et al., 2019). We extracted a subsample of 118 participants (84 males, 34 females, 20 through 35 years old, median = 27.5 years old) for our experiment. The study collected extensive physiological, psychological, and neuroimaging data from healthy adults at the Day Clinic for Cognitive Neurology of the University Clinic Leipzig and the Max Planck Institute for Human and Cognitive and Brain Sciences in Leipzig, Germany. The study followed ethical guidelines outlined by the Declaration of Helsinki and received approval from the ethics committee at the medical faculty of the University of Leipzig. Participants were recruited through various means such as public advertisements, leaflets, online advertisements, and information events at the University of Leipzig. Those who agreed to participate provided written informed consent prior to data acquisition while maintaining anonymity. Monetary compensation was given after completion of all assessment days. Before inclusion in the study’s analysis pool, thorough medical assessments were conducted to ensure control over past or current somatic or mental illnesses as well as medication status.

A 16-minute resting-state EEG was recorded using a BrainAmp MR plus amplifier in an electrically shielded and sound-attenuated EEG booth. The recording used 62-channel active ActiCAP electrodes attached according to the international standard 10–10 localization system, referenced to FCz, with impedance kept below 5 KΩ. The amplitude resolution was set to 0.1 μV, and the bandpass filter ranged from 0.015 Hz to 1 kHz with a sampling rate of 2500 Hz. The session included a total of 16 blocks (8 eyes-closed and 8 eyes-open), where participants fixated their gaze on a black cross presented on a white background during open-eyes segments while seated in front of a computer screen. For the assessment of fluid intelligence, authors employed Subtest 3 of the Performance Testing System (Leistungsprüfsystem 2, LPS-2) which measures logical or inferential thinking. During this subtest, participants are tasked with identifying the symbol in a series that deviates from the logical rule applied to that series, aiming to identify as many of these non-conforming items as possible within a three-minute time frame.

### EEG recording and data preprocessing

All three datasets included the collection of EEG data using 62 or 64-channel devices during the resting state condition. The preprocessing protocol was unified for all the datasets and performed with MNE (Gramfort et al., 2013) and Autoreject libraries (Jas et al., 2017). It included segmentation of the data into epochs of 6 s length with 1 s overlap, filtering them in 1-45 Hz band, re-referencing to an average electrode, resampling the epochs to 100 Hz and processing them in order to remove the artifacts. The artifact removal procedure was fully automated and consisted of ICA-based correction of ocular and muscular artifacts and usage of the Autoreject library to get rid of other artifacts.

#### Source localization and network construction

As a next step, preprocessed data was used to reconstruct the sources of brain electrical activity with the eLORETA algorithm (exact low resolution brain electromagnetic tomography). In the absence of magnetic resonance imaging (MRI) records, we utilized fsaverage object available in the MNE library. To ensure consistency of the obtained results, we performed numerical experiments using two cortical parcellations: Destrieux atlas with 148 regions (74 per hemisphere) and Desikan-Killiany atlas with 68 regions (34 per hemisphere).

The regional time series related to the reconstructed sources were then used to create the functional connectivity matrices. For this end, we utilized three commonly used phase-based functional connectivity (FC) measures: weighted phase lag index (wPLI), phase locking value (PLV) and imaginary part of coherence (imCoh). We applied these methods on the following frequency bands: theta 4-8 Hz, alpha-1 8-10 Hz, alpha-2 10-13 Hz, beta-1 13-20 Hz, beta-2 20-30 Hz, gamma 30-45 Hz.

In formulae below we used the following notation: *E* is a mathematical expectation, *Im* is an imaginary part, *S*_*xx*_ is a power spectral density of a signal and *S*_*xy*_ is a cross-spectral density between two signals.

Weighted phase lag index, a measure of phase lag synchronization proposed by (Vinck et al., 2011) as an improved version of PLI aimed to overcome some of its original disadvantages, such as sensitivity to an uncorrelated noise and volume conduction.

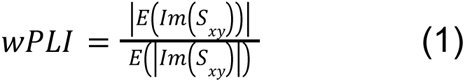

Imaginary part of coherency (imCoh) (Nolte et al., 2004), effectively eliminates potentially spurious instantaneous interactions caused by field spread, by explicitly discarding contributions along the real axis.

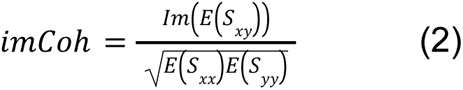

Finally, phase-locking value (PLV) (Lachaux et al., 1999), which represents the absolute value of the mean phase difference between two signals expressed as a complex unit-length vector, was used to estimate degree of synchrony between the reconstructed brain sources.

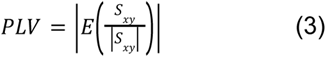

### Connectome-based predictive modeling

Creation of functional connectivity matrices served as a first step for conducting subsequent experiments related to CPM (Shen et al., 2017). In the CPM procedure, a set of functional connectivity matrices and corresponding behavioral data are collected from a specific sample of participants. Each participant contributes an N*N functional connectivity matrix. The CPM procedure operates by iterating through all the edges of these matrices, generating N*(N-1)/2 series of length M, where M refers to the total number of participants. These series are then subjected to a correlation with intelligence score. The edges that exhibit a significant correlation (with a p-value surpassing a predetermined threshold) with the behavioral data are considered valuable. For each participant, the weights of these valuable edges are summed, resulting in a set of M sums. Similarly, a set of M behavioral measures is obtained. These two sets are then used to develop a regression model, which is subsequently utilized for making predictions.

For the default parameter set, the following parameters were chosen: partial correlations (controlling for the effect of age) for correlation analysis, the Destrieux atlas for brain mapping, CPM order of 1, implying that only linear relations are subsumed, and a p-value threshold of 0.001. In addition to this set, numerical experiments were conducted to explore alternative variants. These experiments involved using Desikan-Killiany atlas for brain mapping, and CPM orders of 2 and 3. A p-value threshold of 0.01 was employed as well. Consequently, five main alternative parameter sets were generated, allowing for a comparative analysis against the default one.

### Cross-validation

To assess the performance of each experiment, we utilized a leave-one-out cross-validation (LOOCV) approach. This methodology entailed constructing a predictive model using data from n-1 participants (training set) to predict the score of the remaining participant (test set) in each iteration. The effectiveness of the experiment was assessed by accurately identifying significant edges in at least 95% of the LOOCV iterations.

Experiments that satisfied this criterion were classified as successful, and their outcomes, inclusive of R^2^ (coefficient of determination) and MAE (mean absolute error), were reported. Conversely, experiments that did not meet the specified threshold were deemed unsuccessful and were excluded from the analysis.

### Computational lesioning approach

To clarify the anatomical details of the obtained results, we used computational connectome lesioning, a method that involves systematically removing one important edge at a time during the LOOCV CPM procedure. The difference in the quality of intelligence prediction between the intact and lesioned versions was used to assess the influence of the removed edge. Given the heightened computational demands, we chose not to apply this procedure to all CPM variants but focused instead on the default setting and the variant using the Desikan-Killiany atlas.

### Prediction quality metrics

To assess the quality of prediction we used two metrics, R^2^ and mean absolute error (MAE), that were defined as follows:

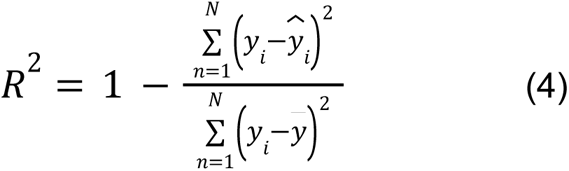

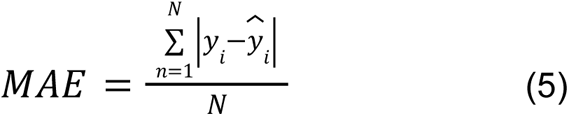

Here *y* is a vector of intelligence values, *ŷ* is a vector of predicted intelligence values, *y̅* is a mean intelligence value and *N* is a sample size. Note that, given this definition, R^2^ can be negative (Scheinost et al., 2019).

We evaluated the statistical significance of the values of these metrics with a permutation test (see below). A result obtained with a specific set of parameters was considered significant if it met two requirements: it was statistically significant (p < 0.05 after multiple comparison correction) and its R² ≥ 0.05.

### Statistical analysis and data visualization

Continuous variables were presented as either mean ± standard deviation for normally distributed data or median (interquartile range, IQR) for data with non-normal distribution. Normality of the distribution was assessed using the Kolmogorov-Smirnov test. Normalization of intelligence values in different datasets was done via implementing a standard min-max normalization procedure. The statistical significance of differences between sex in samples was assessed with a pairwise chi-squared test. Partial correlation analysis was employed to examine the relationships between behavioral and physiological data while controlling for age as a covariate.

The significance of R^2^ values was assessed using a permutation test. Specifically, we randomly permuted observed intelligence values 10,000 times based on predicted intelligence values and calculated the p-value as the proportion of permutations where R^2^ values were equal to or greater than the original case. To address multiple comparisons, individual p-values were corrected using the False Discovery Rate (FDR) method. In the main text of the manuscript we reported only FDR-adjusted p-values. The comparison between whole-brain and lesioned models was performed with Steiger’s z-test (Steiger, 1980). All statistical analyses were performed with scipy (Virtanen et al., 2020) and sklearn (Pedregosa et al., n.d.). Python libraries were used for all calculations, except for the Steiger’s test, for which z-values were calculated using an online calculator (B. A. Weiss, n.d.). Circular plots in the “Default parameter set” and the “Alternative parameter sets” subsections were created with MNE-connectivity visualization tools (Gramfort et al., 2013). Scatter plots in these subsections, as well as categorical plots in the “Lesion results” subsection were generated utilizing the Seaborn library (Waskom, 2021).

## Results

### Demographic and behavioral data of participants

We analyzed data from 3 separate datasets with an aim to predict non-verbal intelligence scores of participants based on resting state EEG data. First, as the intelligence data were normally distributed, we ran ANOVA to compare mean differences in intelligence scores between the datasets. Since all three datasets had different measures of intelligence, the raw data was normalized as described in the “Statistical analysis and data visualization” section prior to the comparison. We discovered a statistically significant main effect of the group (F(2, 252) = 32.63, p < 0.001). Following this, we performed a post-hoc Tukey test to compare datasets pairwise. The results showed that pairs 1–3 and 2–3 differed significantly in terms of mean intelligence scores (both p < 0.001).

Given the non-normal distribution of the participants’ age data, we conducted a Kruskal–Wallis test to evaluate the statistical significance of differences in median values (H = 103.07, p < 0.001). Subsequently, we performed a post-hoc Dunn test for further analysis of these differences. The results indicated a statistically significant difference in medians between sample #1 and #2 as well as between samples #1 and #3 (both p < 0.001).

The statistical significance of differences between sexes in the samples was assessed using a pairwise chi-squared test. The hypothesis of no difference was rejected for samples #1 and #2, and samples #1 and #3 (both p < 0.01). More detailed information about the participants is presented in Table 1. It should be noted that the terms dataset and sample are used interchangeably throughout the text of the article.

**Table 1.** Summary of participants’ demographic and behavioral data.

| Dataset | Number of subjects | Age | Male/female | Intelligence test | Intelligence score |
| --- | --- | --- | --- | --- | --- |
| Dataset # 1 | 72 | 17-25,<br>median = 20<br>(IQR = 2) | 31/41 | Raven's Standard Progressive Matrices | mean = 109.9 (std = 10.8) |
| Dataset # 2 | 65 | 18-35,<br>median = 26<br>(IQR = 10) | 54/11 | Wechsler Adult Intelligence Scale (WAIS-III) | mean = 109.2 (std = 10.8) |
| Dataset # 3 | 118 | 22.5 - 32.5*, median = 27.5 (IQR = 5) | 84/34 | Performance Testing System (LPS-2) | mean = 21.1 (std = 3.4)** |

\*In the 3rd sample age was reported as 5-year intervals, so middle points were taken.
\*\* The intelligence values of the first and second samples were measured using a standard intelligence quotient scale, whereas the third sample values were represented as the number of correct answers.

### Default parameter set

During the analysis of the first dataset, we found significant correlations between intelligence level and functional connectivity in the alpha range. Specifically, in the low alpha band (8–10 Hz), seven edges were selected, which resulted in an R-squared value of 0.12 (p = 0.002, MAE = 7.69). These edges demonstrated a negative correlation with non-verbal intelligence, suggesting that weaker connections were associated with higher intelligence scores. Three main regions, namely the orbital frontal, parietal, and insular cortices, accounted for most of these connections. Additionally, we observed similar associations in the high alpha range (10–13 Hz). Both positively and negatively correlated edges were discovered: 8 edges for positive connections and 18 for negative ones. The model based on positive edges resulted in an R-squared value of 0.11 (p = 0.004, MAE = 7.98), while an R-squared of 0.2 was obtained based on negative edges (p < 0.001, MAE = 7.49; Fig 3). The edges that were positively related to intelligence scores were mainly confined to the occipital, temporal and frontal orbital (polar) cortices. In contrast, negatively correlated links were more numerous and widely distributed over brain areas. All significant associations between brain connectivity and intelligence were obtained using imCoh as a metric.

**Figure 3.**
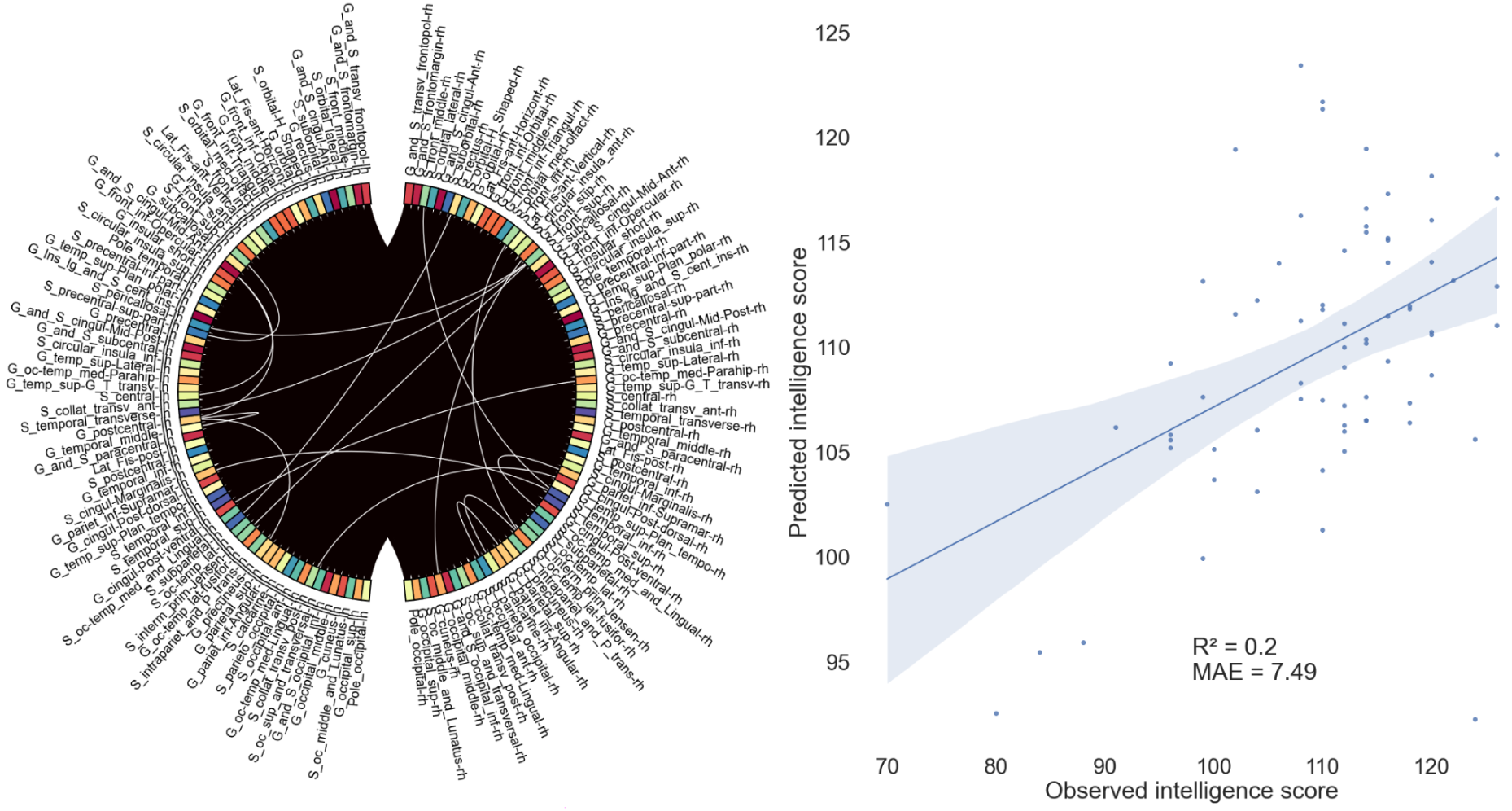
Results of connectome-based modeling for the first sample utilizing the imCoh method in the high alpha range (negatively correlated edges). The left image features a circular plot that illustrates individual brain areas according to the Destrieux parcellation atlas, with functional connections depicted between them. The right image displays predicted intelligence scores plotted against the observed scores.

Similarly, we were able to identify the same relationship between brain and cognitive variables in the second dataset. Five positive edges imCoh-based were located in occipito-parietal regions (R² = 0.09, p = 0.024, MAE = 8.34). The findings from the third dataset were limited to only one significant result, which was found in the theta range (4–8 Hz). Two negative edges in temporo-occipital areas resulted in R² = 0.06 (p = 0.052, MAE = 2.65). In this case, a weighted phase lag index was used to quantify brain connectivity patterns. All the aforementioned associations are summarized in Table 2.

**Table 2.**
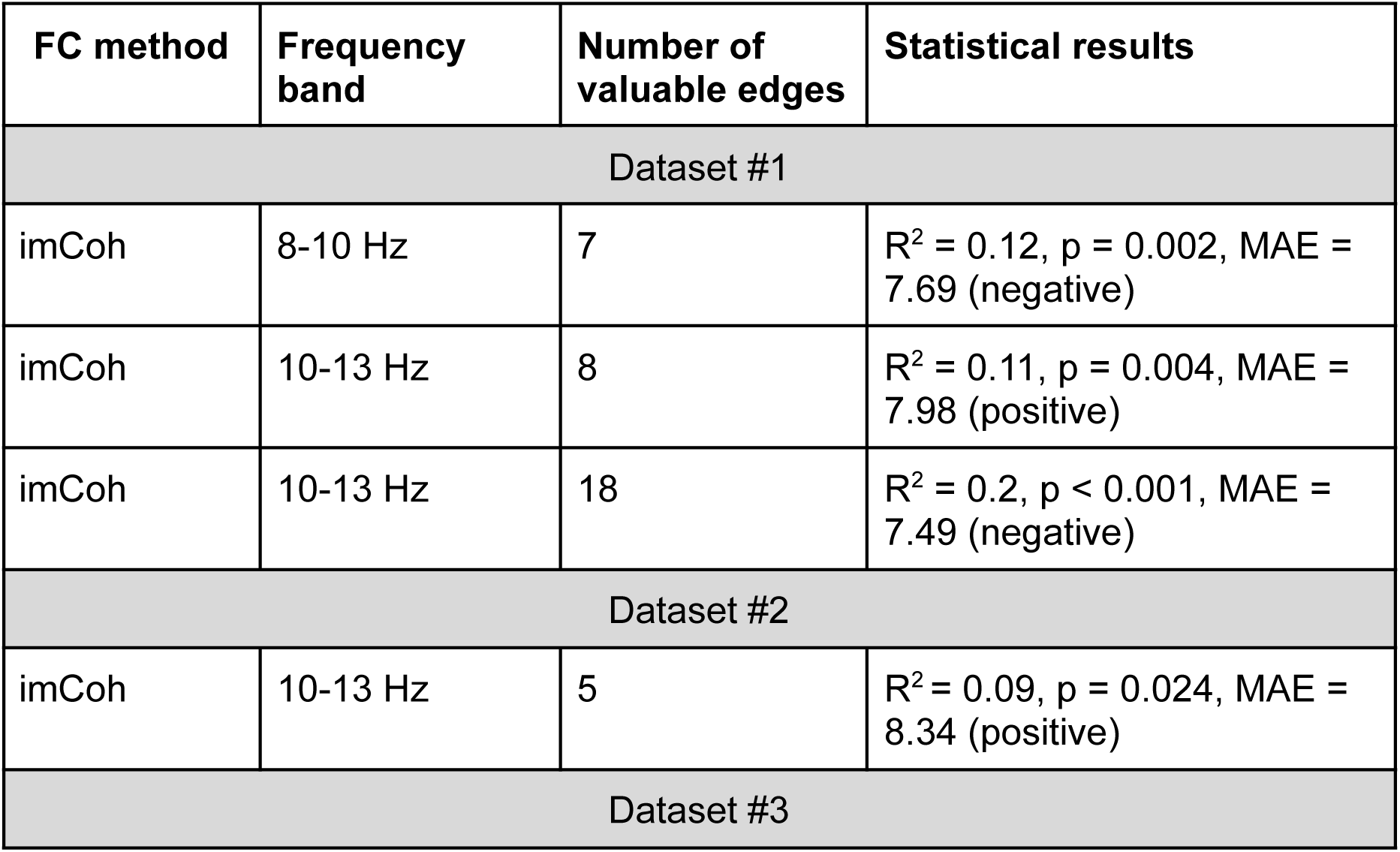
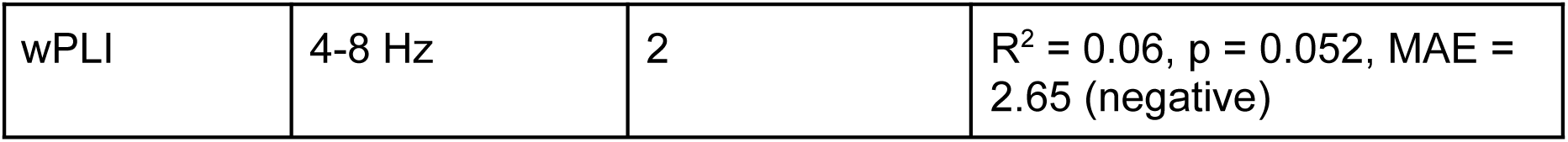
Summary of key results from the CPM analysis utilizing default parameter settings across all three datasets in the study.

## Alternative parameter sets

### p-value threshold

Increasing the p-value threshold by one order of magnitude (to 0.01) yielded the following results. In the first sample, one significant result was observed using the imCoh functional connectivity method for the high alpha band (R² = 0.18, p = 0.001, MAE = 7.64; Fig. 4). Compared to the default variant, the number of significant edges increased to 69, while R² decreased slightly from 0.20 to 0.18. The second sample did not yield any significant results, while the third sample revealed one significant result using phase-locking value in the gamma band with positively correlated edges (R² = 0.05, p = 0.008, MAE = 2.62).

**Figure 4.**
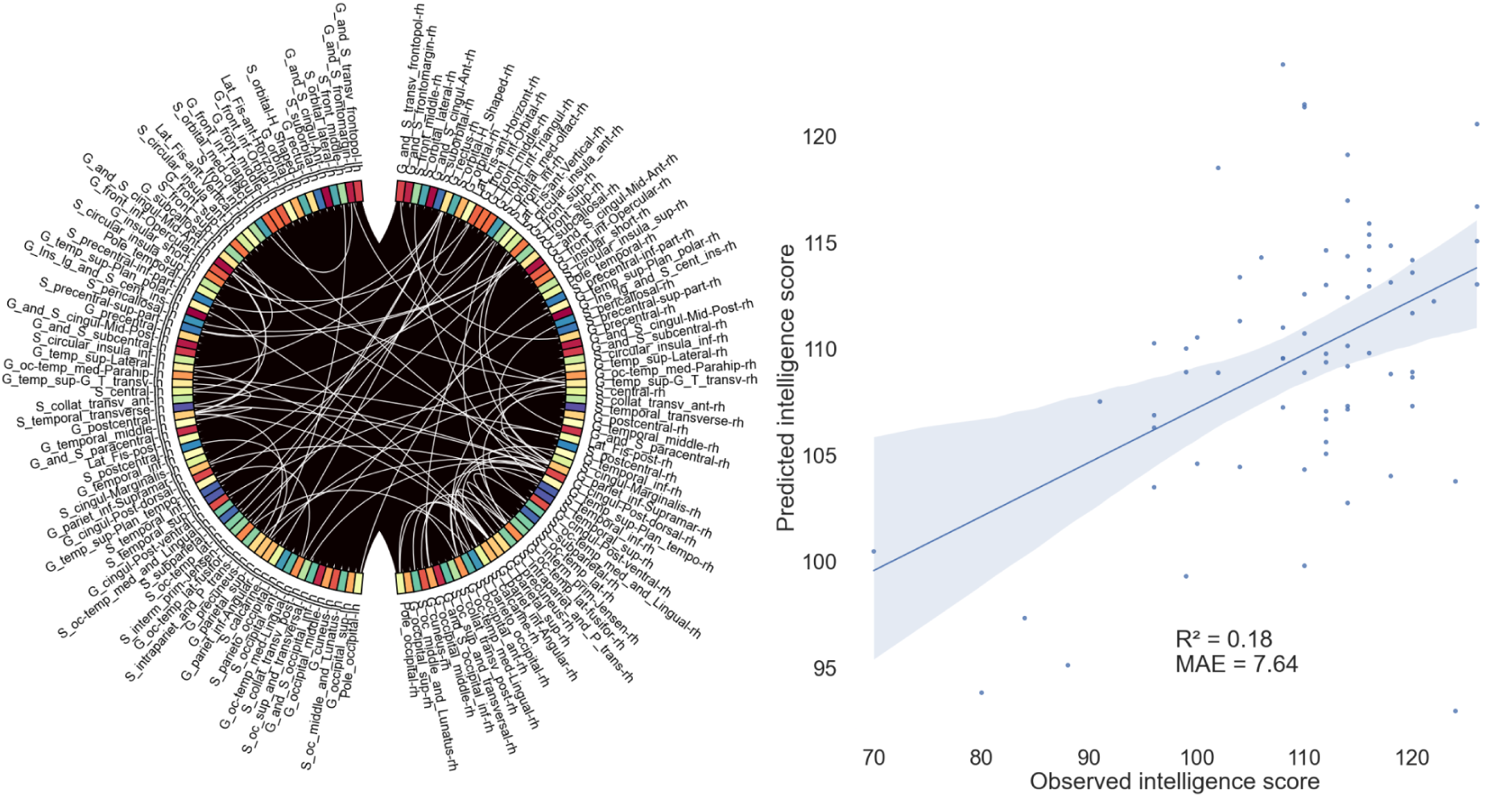
Results of connectome-based modeling for the first sample using the imCoh method in the 10-13 Hz range (negatively correlated edges, p-threshold = 0.01).

### Order of CPM fitting curve

In the analysis of the first sample, three significant results were identified, all derived from the imCoh metric within the alpha frequency band. Specifically, two results were observed in the high alpha band, encompassing both negatively (R^2^ = 0.25, p < 0.001, MAE = 7.0) and positively (R^2^ = 0.12, p = 0.002, MAE = 7.55) correlated edges, while one result was noted in the low alpha band, characterized by negatively correlated edges (R^2^ = 0.1, p = 0.003, MAE = 7.82). The quantity of valuable edges remained constant, as the selection process for edges was unaffected by variations in the CPM order.

When comparing the increased CPM order to the default variant, the outcomes regarding prediction quality were mixed. In one instance, a slight decrease in prediction quality was noted, with the R² value diminishing from 0.11 to 0.12 for imCoh in the 10-13 Hz positively correlated edges. Similarly, a reduction was observed in another case, where the R² value decreased from 0.12 to 0.1 for imCoh in the 8-10 Hz negatively correlated edges. Notably, the optimal performance was achieved with imCoh in the 10-13 Hz negatively correlated edges, where the R² value improved from 0.2 to 0.25 (Figure 5).

**Figure 5.**
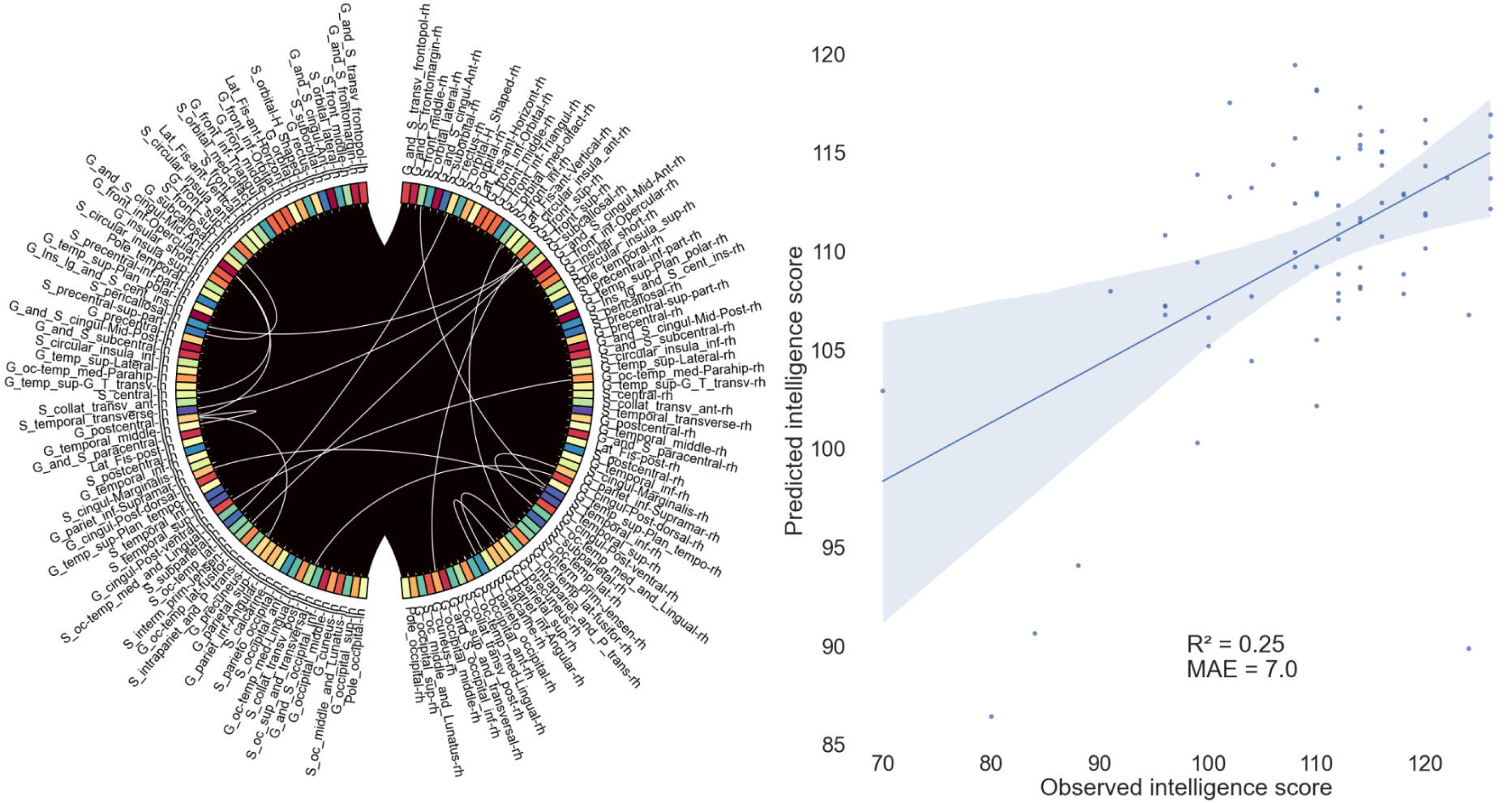
Results of second-order connectome-based modeling for the first sample utilizing the imCoh method in the high alpha range (negatively correlated edges).

The evaluation of the second sample identified one statistically significant result, characterized by imCoh-based positively correlated edges within the high alpha band (R² = 0.08, p = 0.036, MAE = 8.33). This result exhibited a slight decrease in magnitude compared to the default settings. In the third sample, there was also a single significant finding, which involved negatively correlated wPLI-based edges in the theta range (R² = 0.08, p = 0.002, MAE = 2.59). This finding showed a minor increase in magnitude relative to the default analysis, rising from 0.06 to 0.08.

The implementation of third-order CPM did not result in any improvement in predictive accuracy when juxtaposed with second-order models. An examination of the first sample revealed negligible differences compared to the outcomes derived from the second-order CPM. Specifically, within the frequency band of 10-13 Hz, the positively correlated edges exhibited a coefficient of determination of 0.11 (R^2^ = 0.11, p = 0.008, MAE = 7.58), reflecting a minor decline from the 0.12 obtained in the second-order analysis. For the frequency range of 8-10 Hz, the negatively correlated edges sustained an R² of 0.10, indicating stability in this measure (p = 0.006, MAE = 7.76). Furthermore, the negatively correlated edges within the 10-13 Hz band experienced a slight reduction in R² from 0.25 to 0.24 (p = 0.001, MAE = 6.95). In the second sample, the only statistically significant finding aligned with the second-order results, where the positively correlated edge in the 10-13 Hz band demonstrated an R² of 0.05 (p = 0.035, MAE = 8.44). In contrast, the third sample did not yield any significant findings; notably, the previously significant result from the second-order analysis regarding the wPLI-based measures in the 4-8 Hz band, which indicated negatively correlated edges, exhibited a decrease in R² from 0.08 to 0.03, thereby falling below the significance threshold established in the methodology. Table 3 provides a concise summary of the key findings from the analysis conducted.

**Table 3.** Results of comparison of CPM fitting curve orders applied to three datasets.

| FC method | Frequency band | Number of valuable edges | Statistical results | R<sup>2</sup> changes from the default variant (CPM order = 1) |
| --- | --- | --- | --- | --- |
| Dataset #1 (CPM order = 2) |  |  |  |  |
| imCoh | 10-13 Hz | 8 | R<sup>2</sup> = 0.12, p = 0.002, MAE = 7.55 (positive edges) | ↑ (+0.01) |
| imCoh | 10-13 Hz | 18 | R<sup>2</sup> = 0.25, p < 0.001, MAE = 7.0 (negative edges) | ↑ (+0.05) |
| imCoh | 8-10 Hz | 7 | R<sup>2</sup> = 0.1, p = 0.003, MAE = 7.82 (negative) | ↓ (-0.02) |
| Dataset #1 (CPM order = 3) |  |  |  |  |
| imCoh | 10-13 Hz | 8 | R<sup>2</sup> = 0.11, p = 0.008, MAE = 7.58 (positive) | No changes |
| imCoh | 10-13 Hz | 18 | R<sup>2</sup> = 0.24, p = 0.001, MAE = 6.95 (negative) | ↑ (+0.04) |
| imCoh | 8-10 Hz | 7 | R<sup>2</sup> = 0.1, p = 0.006, MAE = 7.76 | ↓ (-0.02) |
| Dataset #2 (CPM order = 2) |  |  |  |  |
| imCoh | 10-13 Hz | 5 | R<sup>2</sup> = 0.08, p = 0.036, MAE = 8.33 (positive) | ↓ (-0.01) |
| Dataset #2 (CPM order = 3) |  |  |  |  |
| imCoh | 10-13 Hz | 5 | R<sup>2</sup> = 0.05, p = 0.035, MAE = 8.44 (positive) | ↓ (-0.04) |
| Dataset #3 (CPM order = 2) |  |  |  |  |
| wPLI | 4-8 Hz | 2 | R<sup>2</sup> = 0.08, p = 0.002, MAE = 2.59 (negative) | ↑ (+0.02) |

### Desikan-Killiany atlas

The first sample yielded four significant findings, consisting of two results derived from wPLI and two from imCoh. The wPLI results were identified within the theta band, revealing positively correlated edges, with one notable edge exhibiting an R² value of 0.06 (p = 0.008, MAE = 8.29). Additionally, a second wPLI result was observed in the high beta band, also indicating positively correlated edges with one significant edge, again yielding an R² of 0.06 (p = 0.008, MAE = 7.53). In contrast, the imCoh results were both assessed within the high alpha band, demonstrating both positively and negatively correlated edges. Notably, the imCoh results exhibited superior R² values compared to those obtained in the default variant, with increases from 0.11 to 0.14 (p = 0.004, MAE = 7.87, four significant edges) and from 0.20 to 0.24 (p < 0.001, MAE = 7.24, four significant edges), respectively. Conversely, the two wPLI results were deemed insignificant in the default variant, and one imCoh result (8-10 Hz, negatively correlated edges, R² = −0.25) was rendered insignificant in this analysis.

The second sample produced five significant results, including four derived from PLV as a functional connectivity method and one from imCoh. The PLV results were identified across various frequency bands: low alpha exhibited positively correlated edges with four significant edges and an R² of 0.27 (p < 0.001, MAE = 7.35; see Figure 6). Concurrently, high alpha showed positively correlated edges with two significant edges (R² = 0.08, p = 0.004, MAE = 8.58). Low beta revealed positively correlated edges with three significant edges (R² = 0.15, p < 0.001, MAE = 8.20). High beta demonstrated positively correlated edges with four significant edges (R² = 0.12, p < 0.001, MAE = 8.04). The sole imCoh result was also identified within the low alpha band, indicating positively correlated edges with two significant edges (R² = 0.26, p < 0.001, MAE = 7.41).

**Figure 6.**
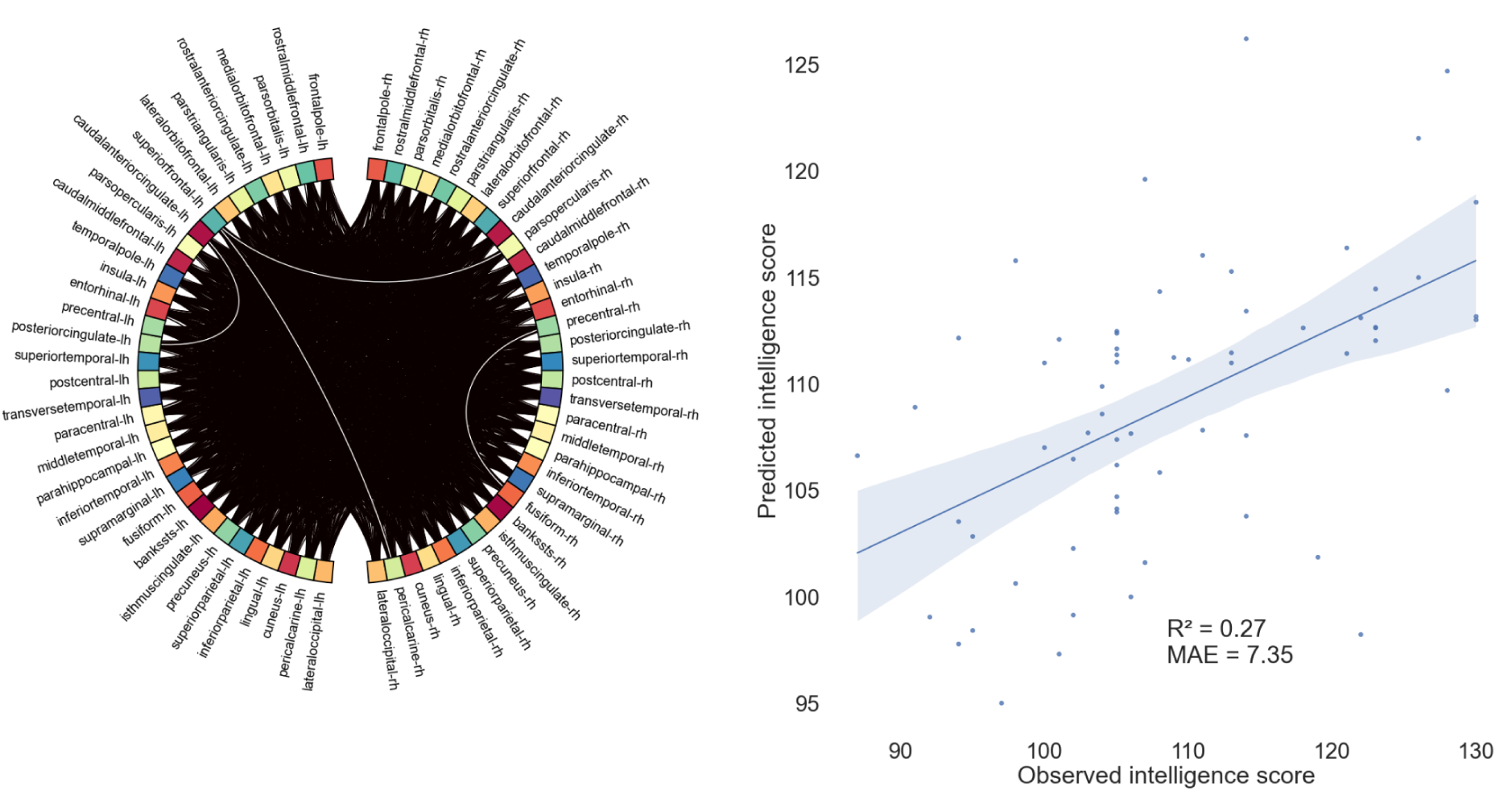
Results of connectome-based modeling for the second sample using the PLV method in the low alpha band range (positively correlated edges).

None of the results demonstrated simultaneous significance when comparing the default variant to the Desikan-Killiany atlas. Notably, the predictive quality of the most favorable outcome improved from 0.09 (imCoh, 10-13 Hz, positively correlated edges) to 0.27 (PLV, 8-10 Hz, positively correlated edges). Furthermore, one result (imCoh, 8-10 Hz, positively correlated edges) exhibited comparability to the default significant finding (imCoh, 10-13 Hz, positively correlated edges) based on the conditions under which it was obtained, specifically utilizing the same functional connectivity method and exhibiting the same sign of correlation in the context of the CPM. However, this result was distinguished by a different frequency band and a lower alpha in the case of the Desikan-Killiany atlas, in contrast to a higher alpha in the default variant. This result also demonstrated enhanced performance, with predictive quality increasing from 0.09 to 0.26. Additionally, the third sample yielded one significant result: imCoh, 20-30 Hz, negatively correlated edges, which included one valuable edge with an R² value of 0.07 (p = 0.0015, MAE = 2.56). This finding cannot be directly compared to the default variant, except in terms of predictive quality, which showed a modest increase from 0.06 to 0.07, as it was derived using a different FC method and within a distinct frequency band. All main results using the DK atlas are presented in Table 4.

**Table 4.**
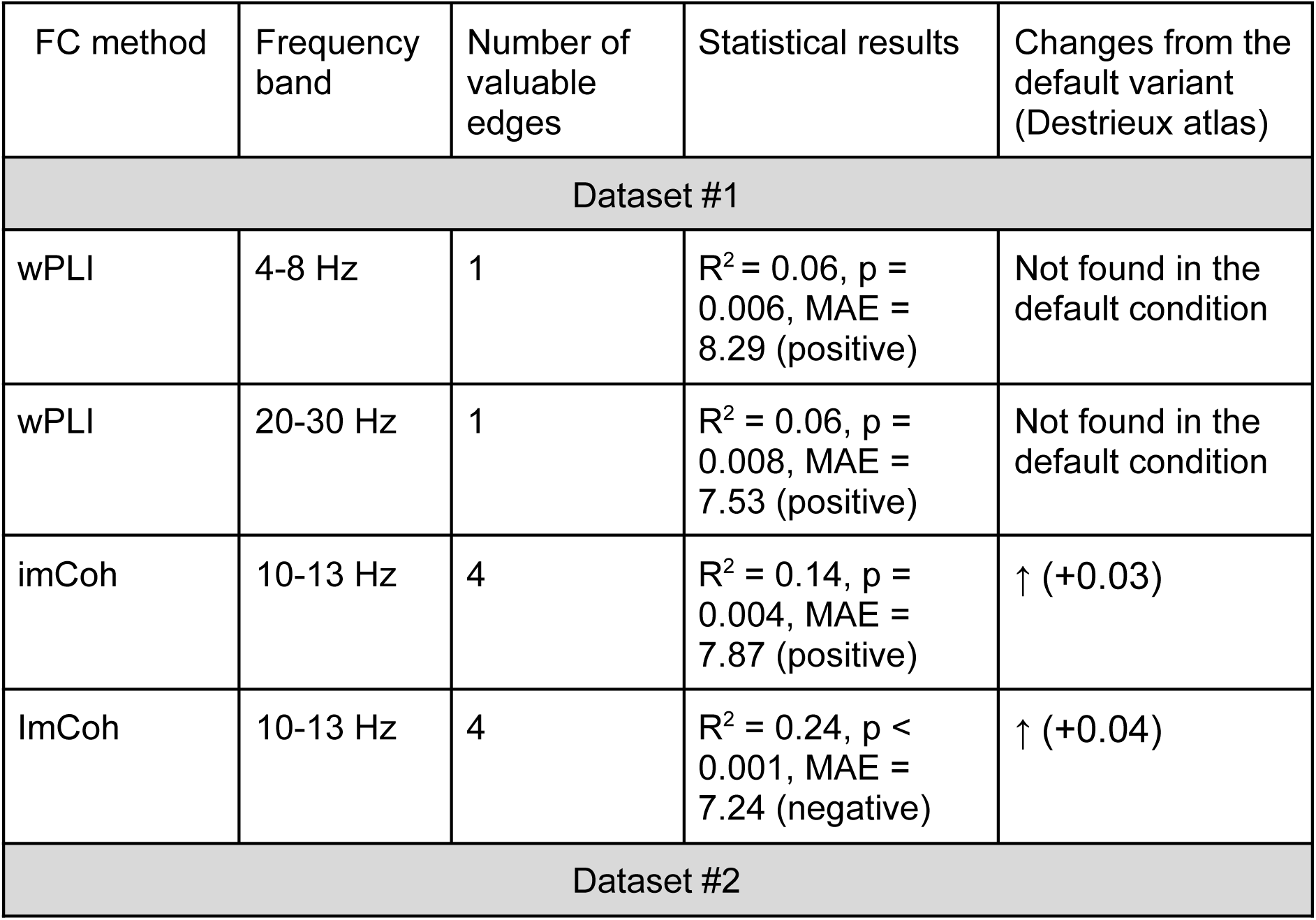
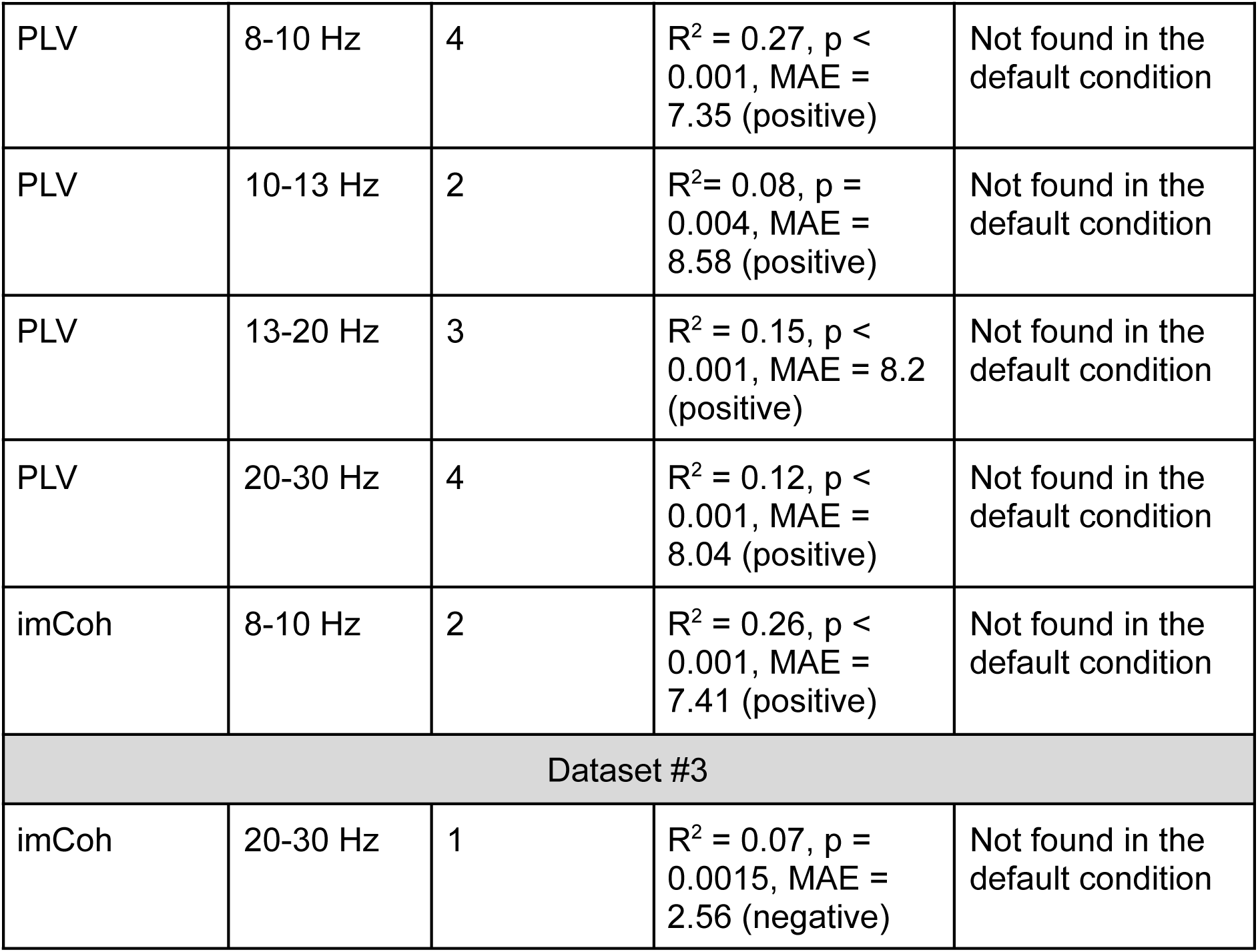
Summary of main results from the CPM analysis employing Desikan-Killiany atlas for cortical regions parcellation.

### External validation

For external validation, we performed the CPM procedure using one sample as a training set and the others as testing sets. The experiment was conducted for all possible pairs of samples. To align the intelligence values of the third sample, which were given on a different scale, we applied the normalization procedure. We used the default parameter set, as well as two of the best alternative variants: the second-order CPM and Desikan–Killiany atlas. Due to the high diversity of samples (different intelligence tests, age, and gender composition), we hypothesized that the external validation would result in low R^2^ values.

Among all the variants, the following combination produced a significant result based on our significance criteria: default parameter set, imCoh FC method, and the frequency band of 20–30 Hz. The third sample was used as the training set, while the second sample served as the testing set. This combination of parameters yielded an R^2^ value of 0.052 (p = 0.33, MAE = 0.2).

### Virtual lesioning approach

In addition to utilizing CPM to forecast intelligence levels based on resting-state EEG data from healthy participants, we aimed to elucidate the impact of specific valuable edges on the overall predictive accuracy by selectively removing or “lesioning” individual edges and recalculating the predictive accuracy of the whole-brain model without that particular connection. Given the computational complexity of these analyses, we limited our investigation to two specific conditions: the default parameter set and transitioning from the Destrieux to the Desikan-Killiany parcellation atlas. To assess the difference between the results of lesioned data and the entire brain, we conducted the Steiger’s z-test, the outcomes of which are detailed in the tables and figures presented below. The Steiger’s test assesses the statistical significance of difference between correlations within two models: the whole-brain model and the lesioned one. These correlations were computed between predicted and observed intelligence values.

### Default parameter set

We investigated three default variants that yielded significant results in the first sample. The analysis demonstrated correlations between 0.34 and 0.37, with corresponding R² values ranging from 0.09 to 0.12, when utilizing imCoh as a method for assessing functional connectivity within the 8-10 Hz frequency band. In the 10-13 Hz frequency band, the correlations increased to between 0.45 and 0.47, with R² values ranging from 0.18 to 0.21. These edges exhibited a negative correlation with intelligence scores. However, these results are not depicted graphically due to the minimal impact of any individual edge deletion on the overall predictive performance of the model.

In contrast, for positively correlated edges within the 10-13 Hz band, correlations were observed between 0.34 and 0.39, with R² values ranging from 0.06 to 0.11 (Fig.7A). The most significant impact on the modeling performance values was observed following the deletion of interhemispheric connection between the orbital part of the right inferior frontal gyrus and left postcentral gyrus (z = 3.2, p = 0.002). The impact on the results was somewhat less pronounced for the removal of intrahemispheric connections between the left orbital part of the inferior frontal gyrus and the superior part of the precentral sulcus (z = 3.00, p = 0.003), as well as between the right transverse frontopolar gyrus and the lateral occipito-temporal (fusiform) gyrus (z = 2.95, p = 0.03).

**Figure 7.**
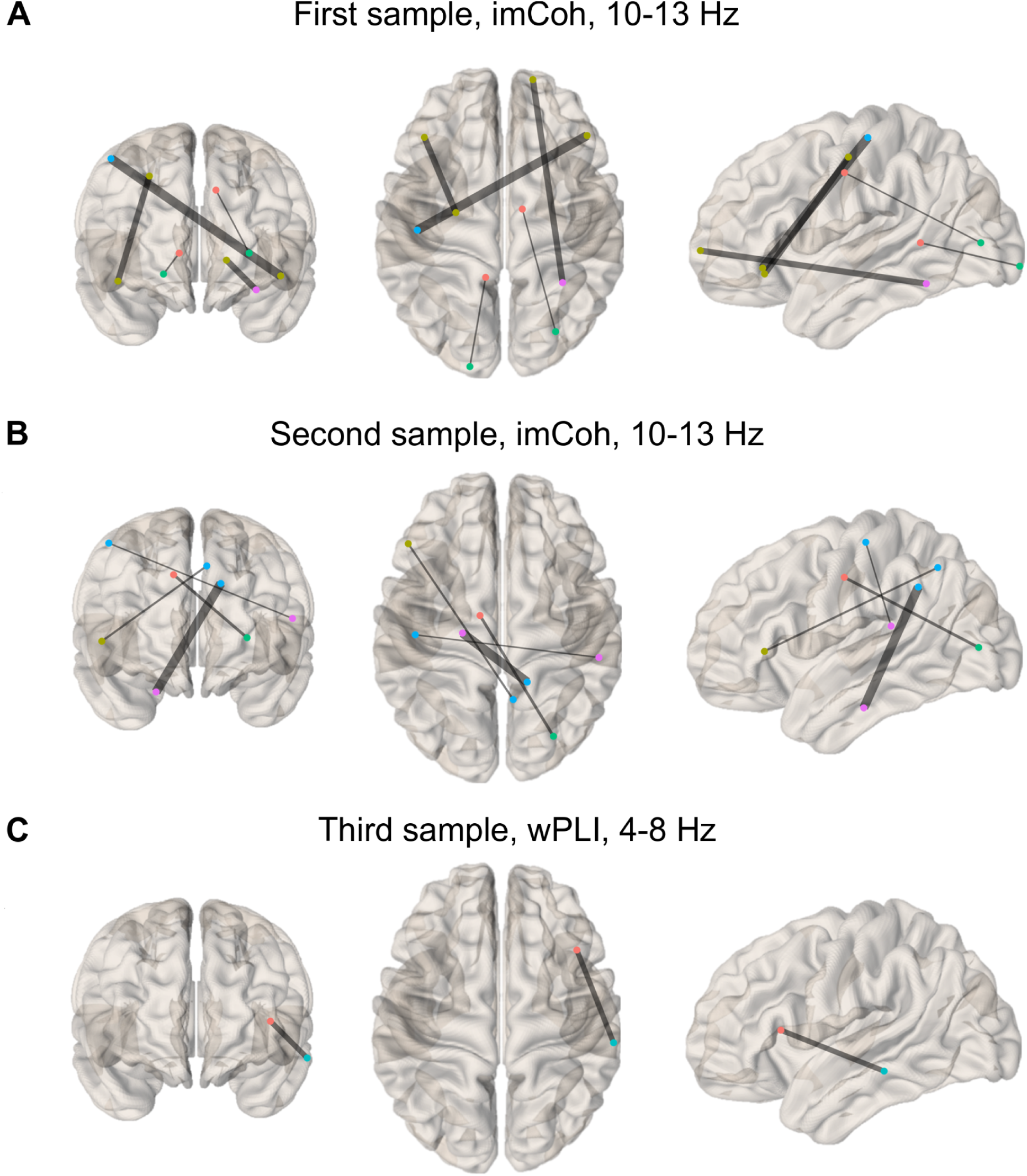
P-values from Steiger’s z-test are presented according to the lesioned edges. The thickness of each edge represents its weight, which is inversely related to the corresponding p-value; thus, a thicker edge indicates a smaller p-value. Panel A displays the first sample (imCoh, 10-13 Hz) with positively correlated edges. Panel B shows the second sample within the same frequency range and FC method. Panel C illustrates the third sample (wPLI, 4-8 Hz) with negatively correlated edges. Node colors represent distinct brain lobes.

The only significant result for the second sample (imCoh, 10-13 Hz, positively correlated edges) indicated that the correlation coefficients ranged from 0.22 to 0.33, with corresponding R² values between 0 and 0.09 (Fig.7B). The most significant influence was observed following the removal of the edge connecting the left medial parahippocampal area to the right subparietal sulcus (z = 3.24, p = 0.001). A slightly lesser influence was detected with the removal of the connection between the left midposterior cingulate gyrus/sulcus and the lunatus sulcus in the right hemisphere (z = 2.84, p = 0.005).

The third sample, which featured wPLI-based valuable edges in the frequency band of 4-8 Hz, revealed significant findings related to negatively correlated edges, with correlation coefficients ranging from 0.14 to 0.28 and R² values ranging from below zero to 0.06 (Fig.7C). In this case, the removal of one of the two existing valuable edges, specifically the intrahemispheric connection between the middle temporal gyrus and the vertical ramus of the lateral sulcus, resulted in a significant decrease in modeling performance (z = 3.02, p = 0.003). Detailed results are illustrated in Figure 9, which includes only those findings exhibiting a minimum difference of 0.05 in correlations between the highest and lowest results for this section. All obtained results and exact correlation values for each edge are presented in Table 5.

**Table 5.** The results of a Steiger’s z-test evaluating the differences between lesioned and whole-brain models. An asterisk sign (*) denotes a significance level of p ≤ 0.05, two asterisks (**) indicate p ≤ 0.01, and three asterisks (***) represent p ≤ 0.001.

| Lesioned edge | Z-value | p-value |
| --- | --- | --- |
| **First sample, imCoh, 10-13 Hz (positively correlated edges)** |  |  |
| Middle-posterior part of the cingulate gyrus and sulcus to middle occipital sulcus and lunatus sulcus | 2.41 | 0.016* |
| Right transverse frontopolar sulcus and gyrus to right lateral occipito-temporal gyrus (fusiform gyrus) | 2.95 | 0.003** |
| Left posterior-ventral cingulate gyrus to left occipital pole | 2.45 | 0.014* |
| Orbital part of the left inferior frontal gyrus to superior part of the left precentral sulcus | 3.00 | 0.003** |
| Orbital part of the right inferior frontal gyrus to left postcentral gyrus | 3.20 | 0.002** |
| Left inferior temporal gyrus to left occipital pole | 1.88 | 0.060 |
| Right inferior temporal gyrus to left calcarine sulcus | 1.15 | 0.251 |
| Right inferior temporal gyrus to anterior segment of the left insular circular sulcus | 1.89 | 0.058 |
| **Second sample, imCoh, 10-13 Hz (positively correlated edges)** |  |  |
| Triangular part of the left inferior frontal gyrus to right precuneus | 2.56 | 0.01* |
| Left parahippocampal gyrus, parahippocampal part of the medial occipito-temporal gyrus to right subparietal sulcus | 3.24 | 0.001*** |
| Right superior parietal lobule to left anterior occipital sulcus and preoccipital notch (temporo-occipital incisure) | 1.27 | 0.21 |
| Left postcentral gyrus to right planum temporale or temporal plane of the superior temporal gyrus | 1.98 | 0.047* |
| Left middle-posterior segment of the cingulate gyrus to right middle occipital sulcus and lunatus sulcus | 2.84 | 0.005** |
| **Third sample, wPLI, 4-8 Hz (negatively correlated edges)** |  |  |
| Right temporal middle gyrus to right vertical ramus of the lateral sulcus | 3.02 | 0.003** |
| Horizontal rami of the anterior segment of anterior segment of lateral sulcus lateral sulcus to left superior occipital and transverse occipital sulcus | 0.97 | 0.33 |

### Desikan-Killiany atlas

In the first sample, out of four significant results identified, two were excluded from further analysis due to the presence of only a single valuable edge. The remaining two variants with imCoh-based valuable edges within the 10-13 Hz frequency band, exhibited both negatively and positively correlated edges. The correlation coefficients for these edges ranged from 0.38 to 0.51 and 0.31 to 0.39, respectively, with corresponding R² values ranging from 0.24 to 0.10 and 0.14 to 0.07 (z = 2.57, p = 0.01 and z = 2.56, p = 0.011). The negatively correlated edge was the connection between the right entorhinal cortex and the parahippocampal area within the same hemisphere. In contrast, the positively correlated edge was the link between the left caudal middle frontal cortex and the right rostral middle frontal cortex (Fig.8A and 8B).

**Figure 8.**
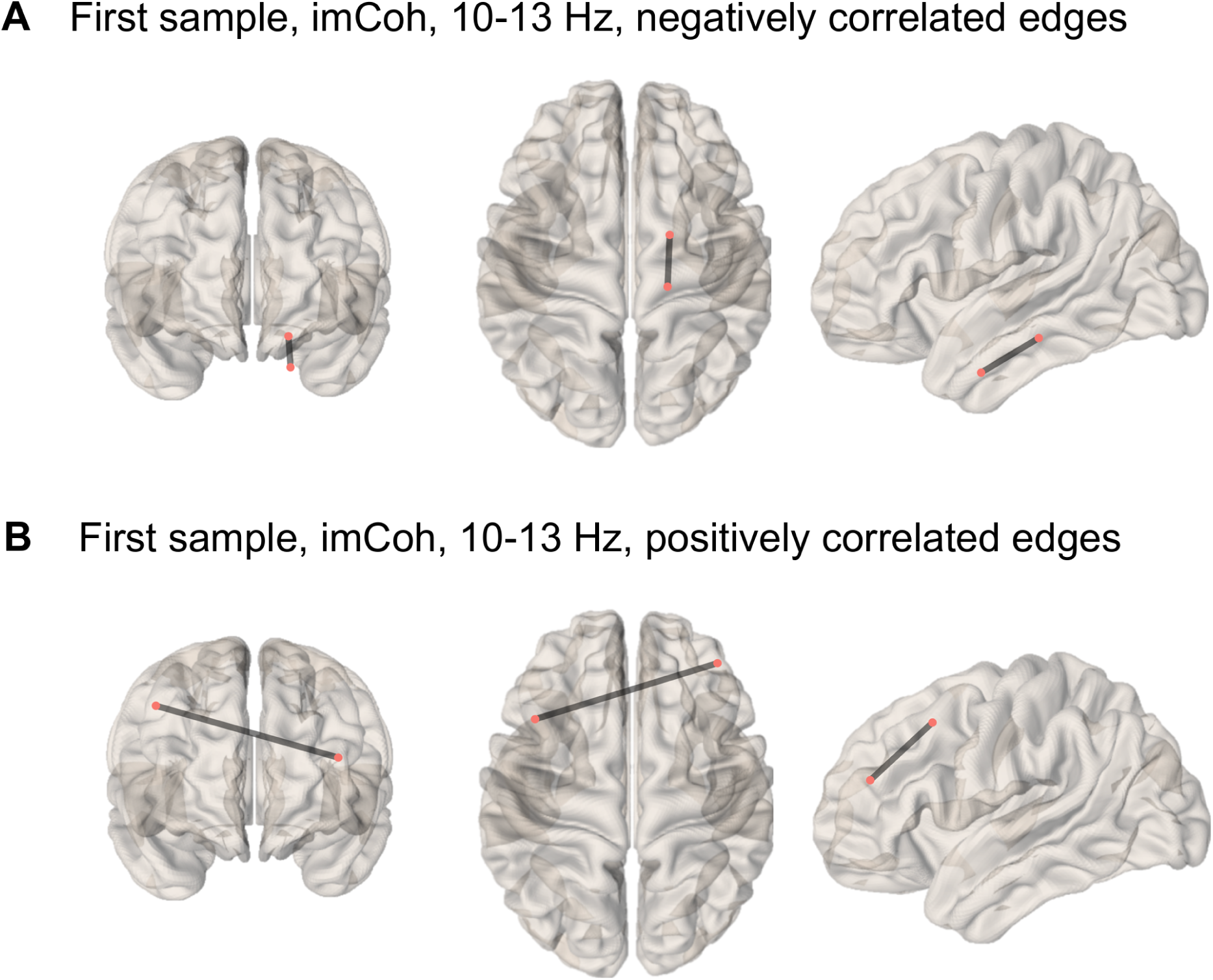
Graphical depiction of Steiger’s z-values derived from the analysis of valuable edge removal in the first sample. Panel A illustrates negatively correlated edges within the imCoh metric at a frequency range of 10-13 Hz. Panel B depicts positively correlated edges under the same conditions.

In the second sample, all results were based on positively correlated edges selected across various frequency bands. The correlation values for these edges were as follows: 0.46 to 0.53 (PLV, 8-10 Hz, R² values from 0.27 to 0.19), 0.15 to 0.34 (PLV, 10-13 Hz, R² values from below zero to 0.07), 0.27 to 0.42 (PLV, 13-20 Hz, R² values from 0.02 to 0.15), 0.32 to 0.39 (PLV, 20-30 Hz, R² values from 0.04 to 0.12), and 0.24 to 0.53 (imCoh, 8-10 Hz, R² values from 0.02 to 0.26). The removal of the connection between the right fusiform gyrus and the right precentral gyrus in the low alpha band significantly affected the modeling results (z = 2.39, p = 0.017). In the high alpha band, two links were identified that significantly influenced the modeling outcomes when removed. Notably, the edge between the right fusiform gyrus and the right precentral gyrus was the most robust, demonstrating a positive correlation with the cognitive measure utilized (z = 3.83, p < 0.001). Furthermore, within the same frequency range, virtual lesioning of the connection between the right pars opercularis and the left superior frontal cortices also had a significant impact on model performance (z = 2.46, p = 0.014; Fig.9A and 9B). Using imCoh as a functional connectivity method, we identified two significant positive edges within the 8-10 Hz frequency range. In this case, left pars orbitalis was connected to the right postcentral (z = 3.02, p = 0.003) and the left posterior cingulate formed a link with the right superior parietal (z = 2.58, p = 0.01; Fig.9C).

**Figure 9.**
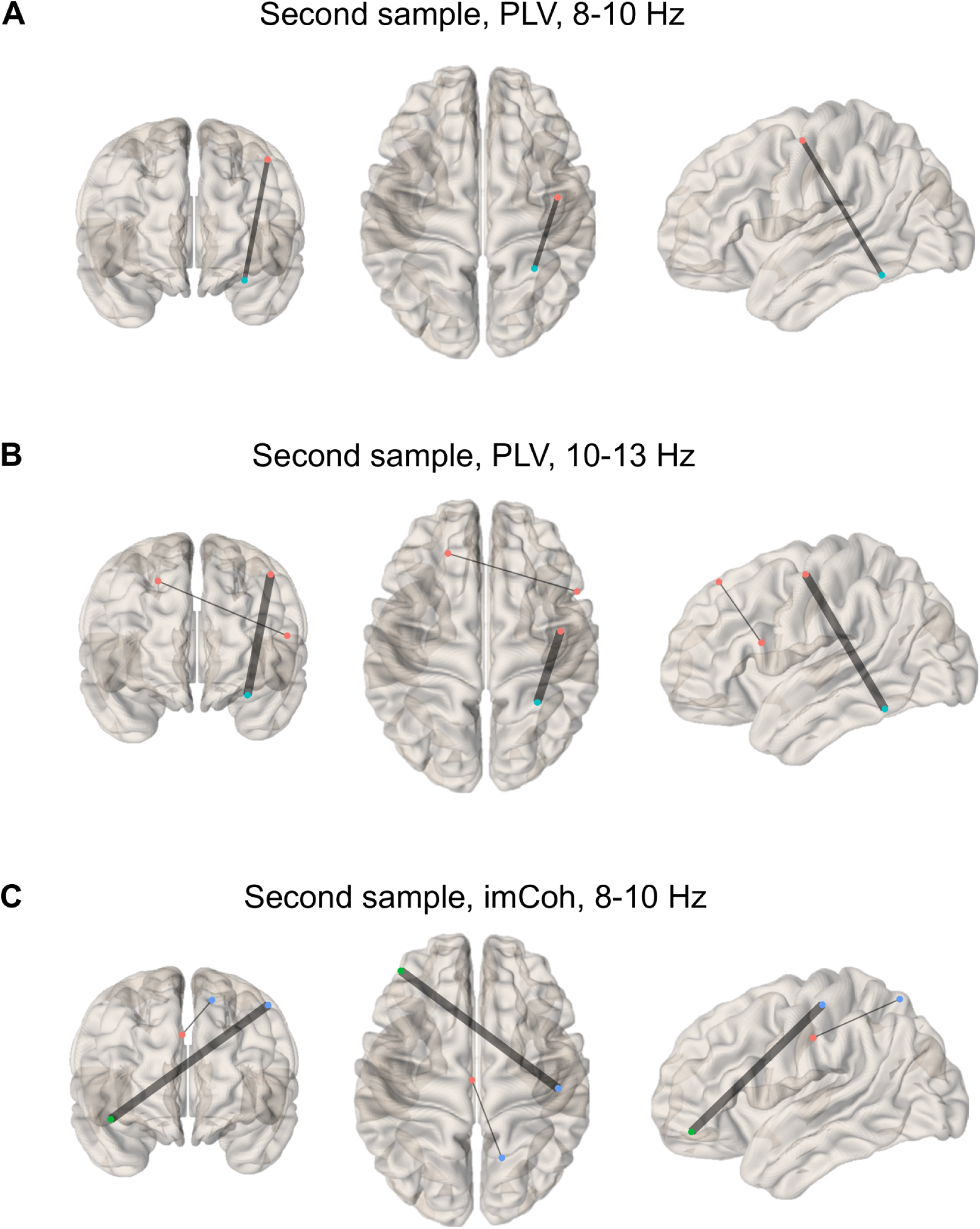
A visual representation of Steiger’s z-values derived from the analysis of valuable edge removal in the second sample (alpha band range). Panel A shows PLV for 8-10 Hz with positively correlated edges. Panel B displays PLV for 10-13 Hz with positively correlated edges. Panel C illustrates imCoh for 8-10 Hz with positively correlated edges.

Similarly, two impactful PLV-based edges were identified in the low beta band: connection from left caudal anterior cingulate to left posterior cingulate cortex (z = 2.65, p = 0.008) and the one linking right fusiform to right precentral gyri (z = 2.42, p = 0.016). Additionally, modeling results in the 20-30 Hz frequency range indicate that temporal and paracentral cortical areas play a crucial role in predicting intelligence, with three noteworthy connections revealed. The edge linking banks of the right superior temporal sulcus to the right postcentral was found to be the most impactful (z = 3.35, p < 0.001). The second most strong intrahemispheric connection bound right fusiform to right precentral (z = 2.94, p = 0.003). Finally, right paracentral to right transverse temporal connection resulted in a drop of prediction accuracy upon removal (z = 2.34, p = 0.019; Fig.10A and 10B).

**Figure 10.**
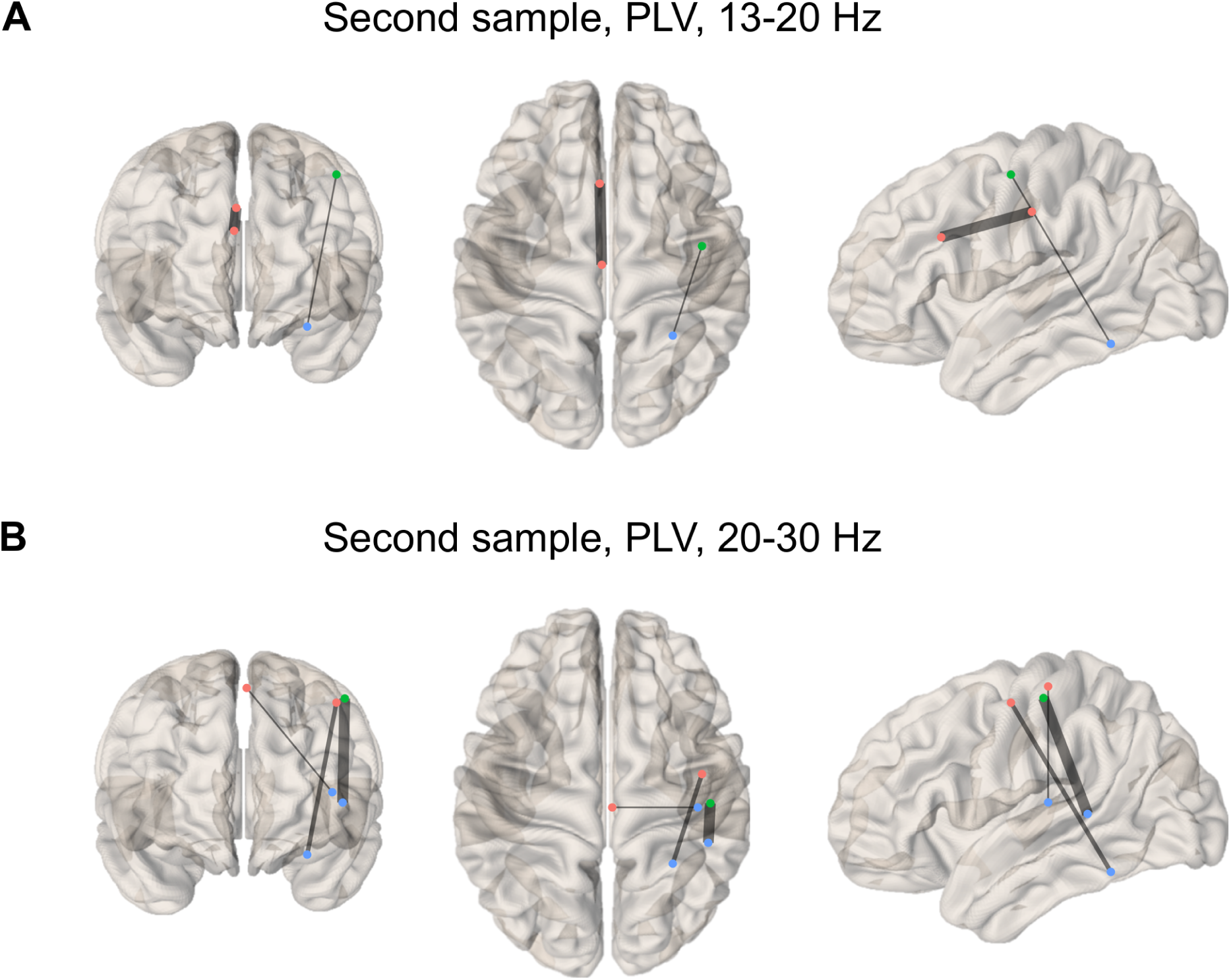
A visual representation of Steiger’s z-values obtained from the analysis of valuable edge removal in the second sample (beta band range). Panel A shows the phase-locking value for the 13-20 Hz range with positively correlated edges, while Panel B displays the PLV for the 20-30 Hz range, also highlighting positively correlated edges.

Notably, no lesion modeling was conducted on the third sample, as its only significant result was associated with a single valuable edge. The main results are summarized in Table 6.

**Table 6.** Results of a Steiger’s z-test assessing the differences between lesioned and whole-brain models based on the Desikan-Killiany atlas.

| Lesioned edge | Z-value | p-value |
| --- | --- | --- |
| **First sample, imCoh, 10-13 Hz (negatively correlated edges)** |  |  |
| Left caudal anterior cingulate to right inferior parietal | 1.01 | 0.31 |
| Right rostral anterior cingulate to right superior orbitofrontal | 0.86 | 0.39 |
| Right entorhinal to right frontal medial orbital | 1.90 | 0.057 |
| Right entorhinal to right parahippocampal | 2.57 | 0.01** |

| **First sample, imCoh, 10-13 Hz (positively correlated edges)** |  |  |
| --- | --- | --- |
| :Left precentral to right rostral anterior cingulate | 1.25 | 0.21 |
| Right precentral to right superior frontal | 1.66 | 0.1 |
| Left caudal middle frontal to right rostral middle frontal | 2.56 | 0.011* |
| Right cuneus to right pars opercularis (inferior frontal) | 0.72 | 0.47 |
| **Second sample, PLV, 8-10 Hz (positively correlated edges)** |  |  |
| Left caudal anterior cingulate to left posterior cingulate | 1.78 | 0.08 |
| Right pars opercularis to left superior frontal | 1.64 | 0.1 |
| Right pericalcarine to left superior frontal | 0.40 | 0.69 |
| Right fusiform to right precentral | 2.39 | 0.017* |
| **Second sample, PLV, 10-13 Hz (positively correlated edges)** |  |  |
| Right pars opercularis to left superior frontal | 2.46 | 0.014* |
| Right fusiform to right precentral | 3.83 | <0.001*** |
| **Second sample, PLV, 13-20 Hz (positively correlated edges)** |  |  |
| Left caudal anterior cingulate to left posterior cingulate | 2.65 | 0.008** |
| Right pars opercularis to left superior frontal | 1.54 | 0.12 |
| Right fusiform to right precentral | 2.42 | 0.016* |
| **Second sample, PLV, 20-30 Hz (positively correlated edges)** |  |  |
| Banks of the right superior temporal sulcus to right postcentral | 3.35 | <0.001*** |
| Right paracentral to right transverse temporal | 2.34 | 0.019* |
| Right pars opercularis to left superior frontal | 1.35 | 0.18 |
| Right fusiform to right precentral | 2.94 | 0.003** |
| **Second sample, imCoh, 8-10 Hz (positively correlated edges)** |  |  |
| Left pars orbitalis to right postcentral | 3.02 | 0.003** |
| Left posterior cingulate to right superior parietal | 2.58 | 0.01** |

## Discussion

### Alpha band and cognitive functioning

Understanding the emergence of cognitive functions from the intricate neural network poses a significant scientific challenge. One promising strategy to address this challenge involves leveraging machine learning tools for the analysis of brain imaging data. In this study, we capitalized on three distinct cohorts of healthy individuals to assess the feasibility and robustness of predicting intelligence levels using resting-state high-density EEG data through the application of a connectome-based predictive modeling approach. To ensure result consistency, we implemented various data processing pipelines. Our findings revealed that the most reliable outcomes, showcasing high predictive accuracy across different functional connectivity methodologies and datasets, were observed in the alpha band frequencies. It has long been established in neuroscience that the neural mechanisms underpinning alpha rhythm generation play a crucial role in establishing and maintaining an excitation-inhibition balance in the brain. This process lays the foundation for the emergence of complex dynamic brain processes, which are essential prerequisites for enabling advanced reasoning capabilities and abstract thinking.

Previous studies conducted over the past decades have consistently emphasized the pivotal role played by alpha oscillations in actively shaping cognitive abilities. Specifically, concerning general intelligence, research has unveiled significant associations between alpha rhythm frequencies and RSPM scores (Anokhin & Vogel, 1996). Individuals with higher IQ levels exhibited reduced activation in the upper alpha band during simple cognitive tasks compared to those with lower IQ scores (Doppelmayr et al., 2005). Moreover, the general intelligence factor has been strongly linked to the individual alpha peak (Grandy et al., 2013). Building on these findings, our recent investigations have indicated that the modulation of alpha rhythm with task difficulty may be linked to processes involved in encoding, retention, and recognition, serving as part of the neural mechanisms governing cognitive effort regulation (Zhozhikashvili et al., 2022). Furthermore, our prior studies have demonstrated a positive correlation between global network integration metrics in the alpha band and non-verbal intelligence, particularly in the more challenging task blocks of the RSPM (Feklicheva et al., 2021). These and numerous other existing studies support the hypothesis that cortical computations mediated by the alpha band are central to the cognitive processes underlying sophisticated cognitive skills and conceptual reasoning.

### Functional connectivity metrics and the predictability of intelligence

In this study, we observed that the highest values of the R² coefficient were associated with the use of imCoh as a measure of functional connectivity, while the utilization of wPLI and PLV led to comparatively poorer predictions. The potential rationale behind this finding may stem from the characteristics of the imaginary part of coherence. The development of this method aimed to mitigate spurious interactions arising from volume conduction (Nolte et al., 2004). It has also been demonstrated that imCoh is resilient to common input connectivity artifacts for similar reasons (Bastos & Schoffelen, 2016). While these attributes are primarily pertinent to the sensor space (with imCoh originally designed to address sensor space issues), the potential for artificial connectivity concerns may also extend to the source space. The possible explanation of imCoh as the most consistent predictor of intelligence is based on the notion that CPM may be susceptible to spurious false connections. If an edge is deemed “truly valuable,” meaning its strength aligns with intelligence levels, it should exhibit consistent changes across the sample as participants’ intelligence varies. Conversely, if edge connectivity is influenced by a common source, its strength will fluctuate based on the neural dynamics of this source, thereby obscuring the relationship with intelligence and diminishing the intelligence-edge weight correlation. If this effect is pronounced, the CPM algorithm is likely to disregard the edge as valuable (given the weakened correlation), consequently compromising prediction accuracy.

An intriguing question arising from this hypothesis is why wPLI failed to yield comparable outcomes despite sharing the property of averting false interactions from a common source (Vinck et al., 2011). Moreover, both wPLI and imCoh have demonstrated similar performance across various applications (Yoshinaga et al., 2020; Zakharov et al., 2020). On the other hand, some variability does exist (Candelaria-Cook & Stephen, 2020; Fraschini et al., 2021). In a pure computational sense, imCoh and wPLI differ significantly in that imCoh can be negative while wPLI is always positive. The relationship between signed wPLI and imCoh is roughly one-to-one, especially near zero (Nolte et al., 2020), but the relationship between unsigned wPLI (commonly used in research) and imCoh is not one-to-one across all intervals symmetric to zero. This discrepancy can lead to misinterpretations. For instance, a positive linear relationship between imCoh and IQ performance in a functional connectivity edge may suggest that higher imCoh values correspond to better IQ performance. However, as imCoh can be negative, this relationship extends to negative values as well, impacting the interpretation. On the other hand, the values of unsigned wPLI do not uniquely correspond to imCoh values due to the lack of sign information, leading to ambiguity in linking wPLI values to IQ. This ambiguity can disrupt the linear relationship between wPLI and IQ, potentially reducing the correlation strength.

### Alternative parameter sets of CPM

An essential aspect of constructing predictive models involves the careful selection of a parcellation atlas, facilitating the delineation of distinct brain regions for computing functional connectivity. Presently, researchers have a diverse array of options for partitioning the entire brain into discrete regions, spanning from a few dozen to over three hundred areas, based on functional connectivity patterns, structural connectivity, cytoarchitecture, myeloarchitecture, gene expression, and receptor distribution (Eickhoff et al., 2018; Zachlod et al., 2023). The accuracy and robustness of predictive outcomes can significantly hinge on the chosen atlas, a decision typically left to the discretion of researchers.

In this study, we evaluated the predictive performance of intelligence levels using two commonly utilized atlases: Desikan-Killiany and Destrieux cortical atlases. A crucial observation concerning cortical atlases within the realm of EEG source reconstruction is that the localization error tends to diminish with increased EEG coverage density (J. Song et al., 2015). This could explain why the Desikan-Killiany atlas, with its more coarse-grained nature, exhibits superior performance, as its regions amalgamate more sources and are potentially less susceptible to localization errors.

The order of the polynomial that fits the behavioral data may affect the prediction quality (Shen et al., 2017) due to the fact that the relation between physiological data and intelligence might not be linear. During the computational experiment, we obtained mostly unchanged results, except for the best one (the 1st sample, imCoh, 10-13 Hz, negatively correlated edges), which showed improved prediction quality (from R² = 0.2 to 0.25) compared to the variant of linear fit CPM. The further increased order did not result in any prediction improvement, which, taking into account virtually no additional computational performance cost, gave evidence to use the 2nd order polynomial to fit the data in the CPM procedure.

Increasing the p-value threshold in the CPM analysis resulted in a decline in prediction quality. Previously significant findings lost their significance as their R-squared values decreased, and even the most accurate prediction experienced a slight performance dip. This outcome can be attributed to the elevated threshold leading to more edges being identified as significant during the edge selection process.The newly selected edges at the heightened threshold displayed higher p-values compared to those under the standard threshold, typically falling within the range of 0.001 to 0.01. Nevertheless, higher p-values often signify weaker correlations, potentially diluting the overall strength of correlation. By incorporating a larger quantity of edges with relatively weak correlations, the composite correlation across all edges may decrease in comparison to the scenario where only highly correlated edges were considered, thereby diminishing the predictive efficacy of the model. This may give rise to a trade-off between excessively low and high p-value thresholds, wherein an optimal value is identified to uphold robust connections for intelligence prediction while filtering out weakly correlated variables. In this study, we compared the thresholds of 0.01 and 0.001, revealing enhanced overall performance with a p-value of 0.001. However, insights from prior research suggest that a p-value of 0.005 could potentially be the optimal threshold choice (Ren et al., 2021).

### Computational lesions and neuroanatomical specifics

To explore the interactions between brain regions that significantly contribute to the overall predictive quality, we implemented a computational lesioning approach. This method involved iteratively excluding one of the significant edges while preserving all others and recalculating the R^2^ coefficient at each step. The obtained data confirm findings from previously published studies and highlight the crucial role of frontal and parietal regions in complex cognitive computations (Anderson & Barbey, 2023; Cipolotti et al., 2023). Furthermore, the results of our study indicate that the precentral and postcentral gyri may also be important areas for executing intricate intellectual processes. Given that our focus in this work was on predicting nonverbal intelligence, the presence of sensorimotor regions in this list is not surprising, as earlier research has repeatedly shown that intelligence relies not only on the activity of associative cortical areas but also on primary/secondary sensory and motor regions of the neocortex (Colom et al., 2009).

The distribution of valuable edges throughout the brain implies that, despite the presence of specific regions influencing the quality of intelligence prediction, intelligence likely operates as a holistic brain phenomenon. Utilizing a global network approach appears to offer a more fruitful description of intelligence compared to a local approach. Recent studies using resting-state fMRI have demonstrated that the correlation between fluid intelligence scores and functional connectivity varies depending on the experimental design and methodologies employed, notably in the DMN, prefrontal cortex, and visual cortices (Lang et al., 2015). State-unspecific patterns of the whole brain were shown to have links with fluid intelligence measured as Penn progressive matrices (Takagi et al., 2019). Additionally, a study related to the CPM paradigm (Wilcox & Barbey, 2023) provided evidence suggesting that global networks explain a significant portion of the variance in fluid intelligence, but the contribution of individual networks is relatively modest in comparison. This research also emphasizes the crucial role played by inter-network connections in this phenomenon. Our findings, combined with these results, lend support to the notion of “distributed intelligence,” which can originate from distinct regions but is not confined to them. Moreover, in this study, although we successfully identified the edges that influence prediction accuracy, there were also instances where the difference between the whole-brain and lesioned models was not substantial, irrespective of the edge removed.

### External validation

During the external validation phase, no statistically significant results with a satisfactory R^2^ value were observed, except for one instance with an R^2^ of 0.052. Several factors may account for this outcome. Firstly, different tests were employed across all three datasets to evaluate nonverbal intelligence levels. Secondly, there were variations in the conditions under which resting-state EEG recordings were conducted in all three datasets. Thirdly, the state of resting wakefulness encompasses a broad spectrum of mental processes among participants, potentially leading to increased variability in brain functional connectivity measures. The sole significant finding should be interpreted with caution due to its relatively low R^2^ value and the discrepancy in parameters compared to the cross-validation cases: significant results in the 20-30 Hz frequency band were only observed with the Desikan-Killiany atlas during cross-validation, whereas the external validation result was obtained using the Destrieux atlas. It’s important to note that this result is considered significant in relation to surpassing the predefined threshold for R², which is set at 0.05. However, it still does not achieve statistical significance, as indicated by a p-value greater than 0.1.

### Comparison with fMRI and general conclusion

Our latest findings contribute additional support for the viability of cognitive assessment, specifically in determining intelligence levels, through the application of connectome-based predictive modeling on resting-state EEG data across diverse datasets. To the best of our knowledge, this study represents the inaugural EEG investigation aimed at quantitatively forecasting nonverbal intelligence levels, while ensuring result consistency across distinct independently procured datasets and data processing pipelines. Prior successful endeavors employing the CPM methodology to predict cognitive performance metrics have been showcased in a succession of recent fMRI inquiries. For instance, Yoo and collaborators, utilizing both resting-state and task-based fMRI, were able to reliably predict attentional capabilities (Yoo et al., 2018). In a related study, researchers compellingly illustrated the CPM approach’s efficacy in forecasting indices of cognitive reserve (Boyle et al., 2023). Our findings suggest the feasibility of achieving comparable predictive accuracies for cognitive abilities using resting-state EEG data as an alternative to relying solely on fMRI data. Nonetheless, it is crucial to underscore the mounting consensus among researchers regarding the advantages of incorporating task-based neuroimaging data rather than analyzing resting-state recordings only. We anticipate that forthcoming studies employing task-oriented designs will address this pertinent consideration.

The methodology employed in this study holds promise for yielding novel insights into the neural underpinnings of intelligence and for advancing the development of more precise and efficient tools for evaluating cognitive functions. Concurrently, this research vividly underscores the impact of methodological choices, such as functional connectivity techniques, parcellation atlases, and various data processing parameters, on the outcomes obtained. The discretion in selecting these elements largely rests with researchers themselves, emphasizing the imperative of meticulous consideration to ensure robust and reproducible findings. By leveraging the CPM approach on resting-state task EEG data, we stand to deepen our comprehension of the brain’s functional organization and its role in shaping individual differences in cognitive capacities. Furthermore, the utility of this methodology extends to diverse domains including education, psychiatry, and neuroscience, where precise and dependable assessments of cognitive functions hold immense significance.

### Limitations

Our study had a number of limitations. The age data was used as a covariate in calculating the partial correlations within the CPM procedure, but the age of participants of the third sample was provided as 5-years intervals instead of precise values, so the middle point of every interval was taken. The first sample contained only young adults aged 17-24 y. o. and therefore differed from two other samples. Another potential limitation of this study is the absence of an evaluation of the influence of participants’ sex on the accuracy of intelligence prediction.

EEG functional connectivity studies encounter the need to decide between constructing connectivity scores from electrode data (sensor space) or reconstructed brain sources (source space). Each approach has its own strengths and weaknesses, and the results in terms of intelligence prediction may emerge solely from the sensor space approach (Zakharov et al., 2020) or both approaches (Langer et al., 2012), depending on the experimental design.

EEG-based investigations often encounter challenges in studying subcortical structures, as distinguishing subcortical activity from cortical activity can be a significant issue due to the weaker signal from subcortical structures, which are obscured by cortical activity. In light of this limitation, our study concentrated exclusively on cortical structures, despite existing evidence suggesting the involvement of subcortical structures in intelligence, both structurally (Basten et al., 2015; Burgaleta et al., 2014; Grazioplene et al., 2014; Santarnecchi et al., 2017) and functionally (M. Song et al., 2008).

Another factor that could limit the generalizability of the findings is the variety of intelligence tests employed in the study. Each sample underwent a different test, and although all tests aimed to evaluate non-verbal intelligence, each test might involve distinct neurophysiological mechanisms, which could result in them not being interchangeable in terms of functional connectivity interpretations.

Finally, the number of valuable edges prior to the virtual lesioning procedure varied across different models.Therefore, future studies should thoroughly investigate the potential impact of this factor on experimental outcomes.

## Abbreviations

EEG: electroencephalography
fMRI: functional magnetic resonance imaging
CPM: connectome-based predictive modeling
PLV: phase locking value
wPLI: weighted phase lag index
imCoh: imaginary part of coherence
IQ: intelligence quotient
RSPM: Raven’s standard progressive matrices
LOOCV: leave-one-out cross-validation
MAE: mean absolute error

## Data availability statement

Datasets #2 and #3 are available at https://chbmp-open.loris.ca/ and http://fcon_1000.projects.nitrc.org/indi/retro/MPI_LEMON.html, respectively. Dataset #1 is available from the corresponding author upon reasonable request. The code used during the processing pipeline is openly available at https://github.com/Goliath-dev/IntelligenceCPM.

## Notes

https://chbmp-open.loris.ca/

http://fcon_1000.projects.nitrc.org/indi/retro/MPI_LEMON.html

https://github.com/Goliath-dev/IntelligenceCPM

### Competing Interest Statement

The authors have declared no competing interest.

